# Decoding Prokaryotic Whole Genomes with a Product-Contextualized Large Language Model

**DOI:** 10.64898/2025.12.03.692003

**Authors:** Shiwen Ni, Shuaimin Li, Shijian Wang, Wenzhi Xue, Luoxi Zhang, Liyang Fan, Xinping Bi, Yitai Li, Chengguang Gan, Jiarui Jin, Yuan Lu, Ahmadreza Argha, Hamid Alinejad-Rokny, Tong Si, Min Yang, Teng Wang

## Abstract

Genomes encode the instructions for life, yet their interpretation requires models capable of capturing long-range genome-wide functional context at scale. Existing genomic foundation models have primarily focused on sequence-level representations, providing powerful tools for biological prediction while leaving complementary opportunities for function-level genome modeling. Here, we present GenSyntax, a function-level genome representation learning framework that models prokaryotic genomes as ordered sequences of gene-product descriptors. By treating gene products as semantic units, GenSyntax transforms annotated replicons into interpretable “genetic paragraphs” that capture genome-wide functional context. We trained GenSyntax on 49,250 annotated prokaryotic genomes and evaluated its ability across genome-scale tasks, including plasmid host prediction, gene-product disambiguation, genome contig ordering and gene essentiality prediction. Compared with general-purpose large language models and representative genomic foundation models, GenSyntax achieved competitive performance across multiple benchmarks, with independent external evaluations supporting its generalization beyond the original training datasets. GenSyntax embeddings also captured signals associated with microbial phenotypes, and gene essentiality predictions enabled exploratory design of minimal genomes. Together with the lightweight GenSyntax-Tiny model, GenSyntax provides a complementary, post-annotation framework for function-level analysis of prokaryotic genomes.

## Introduction

Genomes encode the fundamental blueprints of life, with nucleotide sequences directing biological complexity through hierarchical patterns that resemble the syntax of human language^1^. Although the genomic “grammar” governing the organization and interactions of genetic elements is not fully understood, recent advances in genome language models have begun to reveal these latent structures^2^. Inspired by large language models (LLMs) in natural language processing (NLP)^2–6^, these genome language models have successfully learned representations from nucleotides, proteins, and other biological features, demonstrating remarkable capabilities across tasks ranging from functional prediction to sequence generation^1,7–19^. Collectively, these approaches establish representation learning as a powerful framework for understanding biological systems across multiple scales.

A central challenge in genome representation learning is how to model genomic information at biologically meaningful scales. Prokaryotic replicons typically comprise thousands of genes organized into coordinated functional systems, whereas most existing genomic foundation models represent genomes through sequence-level or sequence-derived features, such as nucleotides, amino acids or learned biological embeddings^7,12–21^. These representations excel at capturing sequence patterns but are less directly aligned with the functional concepts through which biologists typically interpret genomes. In addition, genome-scale sequence modeling remains computationally demanding despite continued advances in long-context architectures and token-compression strategies^1,2^. These considerations motivate complementary representations that directly encode genome-wide functional organization.

Here we introduce a framework for function-level genome representation learning, in which genomes are represented as ordered sequences of gene product descriptors. This abstraction transforms each replicon into a “genetic paragraph”, where each lexical unit corresponds to a human-interpretable functional annotation rather than a nucleotide or protein sequence^22^. Representing genomes at the level of functional semantics offers several conceptual advantages. It compresses millions of nucleotides into thousands of biologically meaningful tokens, enabling efficient modeling of genome-scale functional context for most prokaryotic replicons. It also improves interpretability by directly linking model representations to functional annotations familiar to biologists, while naturally reflecting genome organization in terms of genes, pathways and functional modules. Importantly, this representation enables models to reason directly over the functional architecture of genomes, including the organization, and contextual relationships among genes and genomic elements. Rather than replacing sequence-based genomic foundation models, this functional abstraction provides a complementary semantic layer for genome-scale functional reasoning.

Building on this representation, we developed GenSyntax, a gene product-contextualized language model for function-level analysis of prokaryotic genomes. Using RefSeq, we curated a corpus of 49,250 annotated prokaryotic genomes and organized their gene-product descriptors into replicon-specific ‘genetic paragraphs’^23^. These functional sequences were used for continuous pre-training of a LLaMA3.1-8B foundation model, followed by task-specific fine-tuning^4,6^. We evaluated GenSyntax across four genome-scale tasks, plasmid host identification, gene-product disambiguation, genome contig ordering, and gene essentiality prediction. Compared with general-purpose LLMs, contemporary genome language models and task-specific methods where applicable, GenSyntax achieved competitive performance across all tasks, with independent external benchmarks further supporting its generalization. In addition, GenSyntax embeddings captured microbial phenotype-associated signals, and its gene essentiality predictions enabled exploratory derivation of candidate minimal genomes. Together with the lightweight GenSyntax-Tiny model and a web interface, GenSyntax provides a complementary framework for function-level, post-annotation modeling of prokaryotic genomes.

## Results

### Development of a gene product-contextualized LLM

We assembled a comprehensive dataset of prokaryotic genomes using all fully sequenced bacterial and archaeal assemblies available in the NCBI RefSeq database as of April 18, 2025^24^. To ensure consistency in modeling replicon-level functional organization, we restricted our dataset to complete genomes with uninterrupted chromosomal and plasmid sequences. This curation yielded 49,250 high-quality genomes, including 48,641 bacterial and 609 archaeal genomes, spanning 63 phyla, 2,273 genera and 12,706 species (Supplementary Fig. S1a).

Each genome in RefSeq was functionally annotated using the NCBI Prokaryotic Genome Annotation Pipeline (PGAP), which identifies protein-coding genes as well as non-coding RNA (ncRNA) elements, including transfer RNAs (tRNAs), ribosomal RNAs (rRNAs), and transfer-messenger RNAs (tmRNAs)^22^. From these annotations, we extracted gene product descriptors, such as “glutathione-disulfide reductase” or “ABC transporter permease”, which describe the annotated functions of genetic elements (Fig. 1a, see Methods for details). This process generated a large corpus of over 201 million descriptors across 73,136 unique gene products. Protein-coding genes accounted for the vast majority (97.75%) of products, followed by tRNA (1.67%) and rRNA (0.41%). In this corpus, 10.13% and 21.66% genes are annotated as “hypothetical protein” in bacterial and archaeal genomes, respectively, while this fraction raises to 26.19% in all plasmids. The frequency-rank distribution of gene products approximately followed a power-law pattern (Supplementary Fig. S1b), showing a Zipf-like distribution consistent with the analogy between genomic organization and natural language structure^25^.

**Figure 1.**
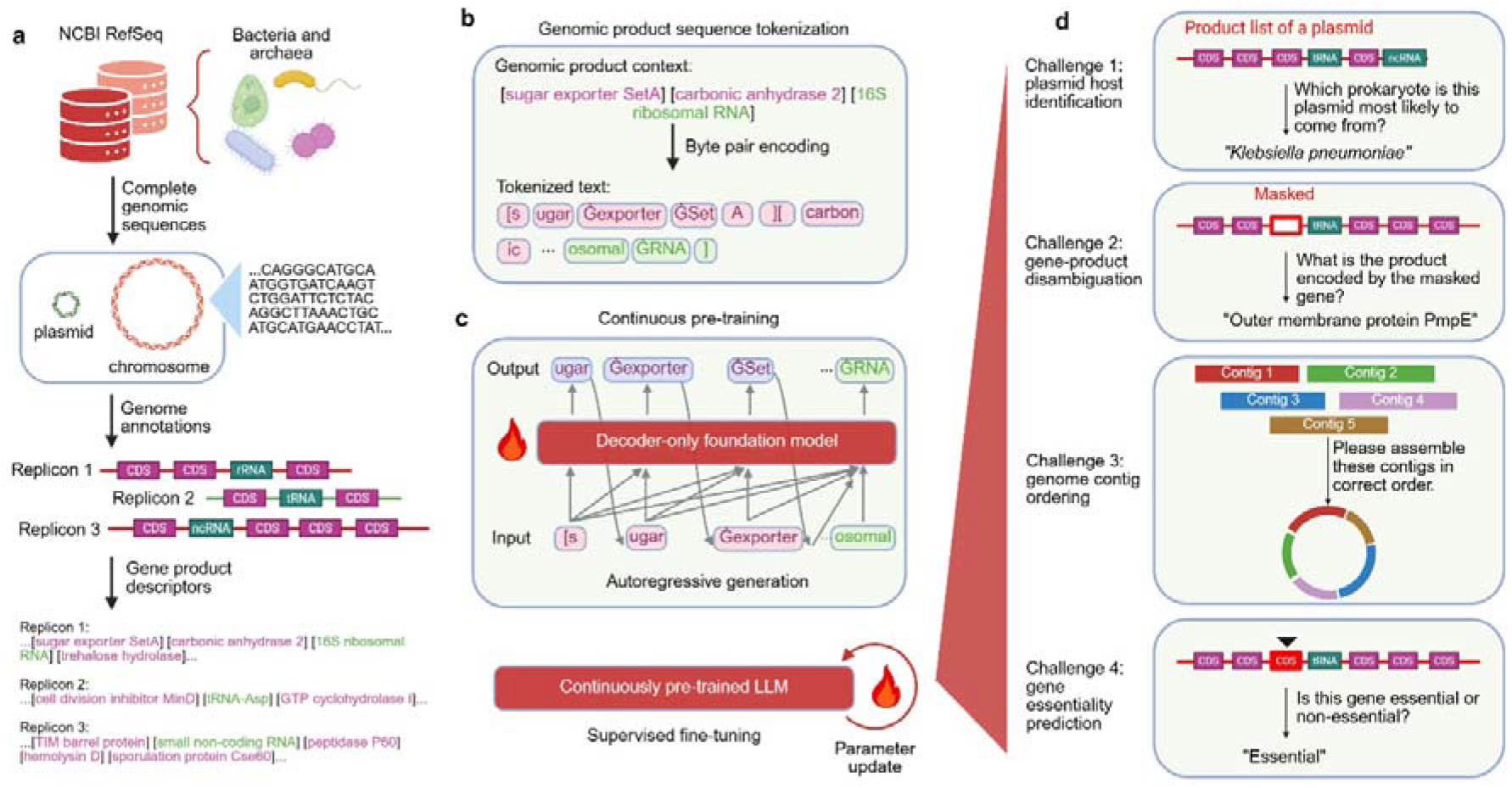
Development and applications of GenSyntax, a gene product-contextualized large language model. **(a)** Construction of prokaryotic genomic semantics through gene-centric linguistic abstraction based on genome annotations. Complete bacterial and archaeal genomes with functional annotations were retrieved from NCBI RefSeq and split into individual replicons (chromosomes or plasmids). For each replicon, gene product descriptors were extracted to generate a functional narrative that encodes genomic context. **(b)** Genomic product contexts were tokenized by byte-pair encoding. **(c)** Tokenized genomic contexts were used to continuously pre-train the base LLM, a decoder-only foundation model, by autoregressive generation. **(d)** The pre-trained model was fine-tuned on four biologically relevant tasks: (1) plasmid host prediction based on encoded product profiles; (2) prediction of masked gene products from replicon product lists; (3) context-aware ordering for genome contigs; and (4) gene essentiality prediction. Model parameters were continuously updated during fine-tuning.

For genomes comprising multiple replicons, we partitioned them into discrete “paragraphs”, each corresponding to an individual replicon (chromosome or plasmid). Within each replicon, gene product descriptors were ordered according to their genomic coordinates, resulting in 51,828 chromosomal and 60,758 plasmid paragraphs (Fig. 1a, Supplementary Figure S1c-e). This transformation enabled annotated replicons to be represented as structured sequences of human-readable functional descriptors, achieving substantial dimensionality reduction while retaining functional context.

Using this corpus, we developed GenSyntax through a two-stage training strategy. In the first stage, we continuously pre-trained the base LLaMA3.1-8B model on chromosomal and plasmid paragraphs to learn general genomic semantics (Fig. 1b and c)^6^. In the second stage, we fine-tuned the model using domain-specific instruction datasets, also derived from the gene product corpus, targeting four biologically relevant challenges: plasmid host identification, gene-product disambiguation, genome contig ordering and gene essentiality classification (Fig. 1d). In parallel, we generated GenSyntax-Tiny by full-parameter training based on Qwen3-0.6B on the same gene product corpus, using the same training objective and data construction strategy^26^. The main GenSyntax training was performed on a cluster of 40 NVIDIA H800 GPUs (80 GB each) over a period of approximately one month. Across all context-specific tasks, we benchmarked GenSyntax against representative genomic foundation models spanning different representation paradigms, including nucleotide-based models (NT-v2, Evo2 and gLM2) and protein-centric models (Bacformer)^12–15,18^. While gLM2 is mixed-modal, we restricted our use to its nucleotide mode. Although these models operate on different biological modalities, all were evaluated using the same downstream protocol.

### Context-specific challenge 1: Plasmid host identification

Plasmids represent a major class of extrachromosomal genetic elements in prokaryotes, frequently carrying genes that confer adaptive traits such as antibiotic resistance, metabolic versatility, and pathogenicity^27^. Natural environments exhibit vast plasmid diversity, as evidenced by resources like the IMG/PR database, which catalogs over 700,000 plasmids^28^. However, for the majority of plasmids, particularly those recovered from metagenomic samples, their host organisms remain unknown^29^. Resolving these host-plasmid relationships is critical for understanding microbial ecology, horizontal gene transfer, and the dissemination of resistance genes. Traditional approaches to plasmid-host prediction typically rely on sequence-based features such as *k*-mer composition or replicon typing^30–33^. Here, we asked whether a function-level genome representation based on gene-product descriptors could capture additional functional-context signals linking plasmids to their host taxa.

We curated a dataset of 60,613 prokaryotic plasmids, each encoding at least four annotated genes and linked to known host taxonomic labels. Of these, 59,613 plasmids were used to fine-tune GenSyntax, enabling the model to associate functional content with host identity across multiple taxonomic levels, from order to strain (Fig. 1d). The remaining 1,000 plasmids were held out as an independent test set to evaluate performance on previously unseen examples (Supplementary Fig. S2). Plasmids in the test set were completely excluded from pretraining, ensuring that the model was not evaluated on previously encountered plasmid sequences. We further characterized the taxonomic composition of the test set and confirmed its consistency with the overall dataset distribution (Supplementary Fig. S3).

We benchmarked GenSyntax against general-purpose LLMs, contemporary genomic foundation models including NT-v2, gLM2, Evo2 and Bacformer^12–15,18^, as well as the specialized plasmid-host prediction tool HOTSPOT^34^. GenSyntax achieved strong host prediction performance across taxonomic levels, reaching accuracies of 0.94, 0.91, 0.79, 0.62 and 0.24 for order, family, genus, species and strain prediction, respectively (Fig. 2a), with consistently strong precision, recall and F1 scores (Supplementary Fig. S4). Compared with general-purpose LLMs, GenSyntax showed consistently improved performance across all taxonomic ranks. Compared with genomic foundation models or HOTSPOT, GenSyntax achieved competitive or superior performance at all taxonomic levels. The lightweight GenSyntax-Tiny model retained comparable predictive performance, indicating that functional gene-product representations remain informative even with reduced model capacity (Fig. 2a and Supplementary Fig. S4).

**Figure 2.**
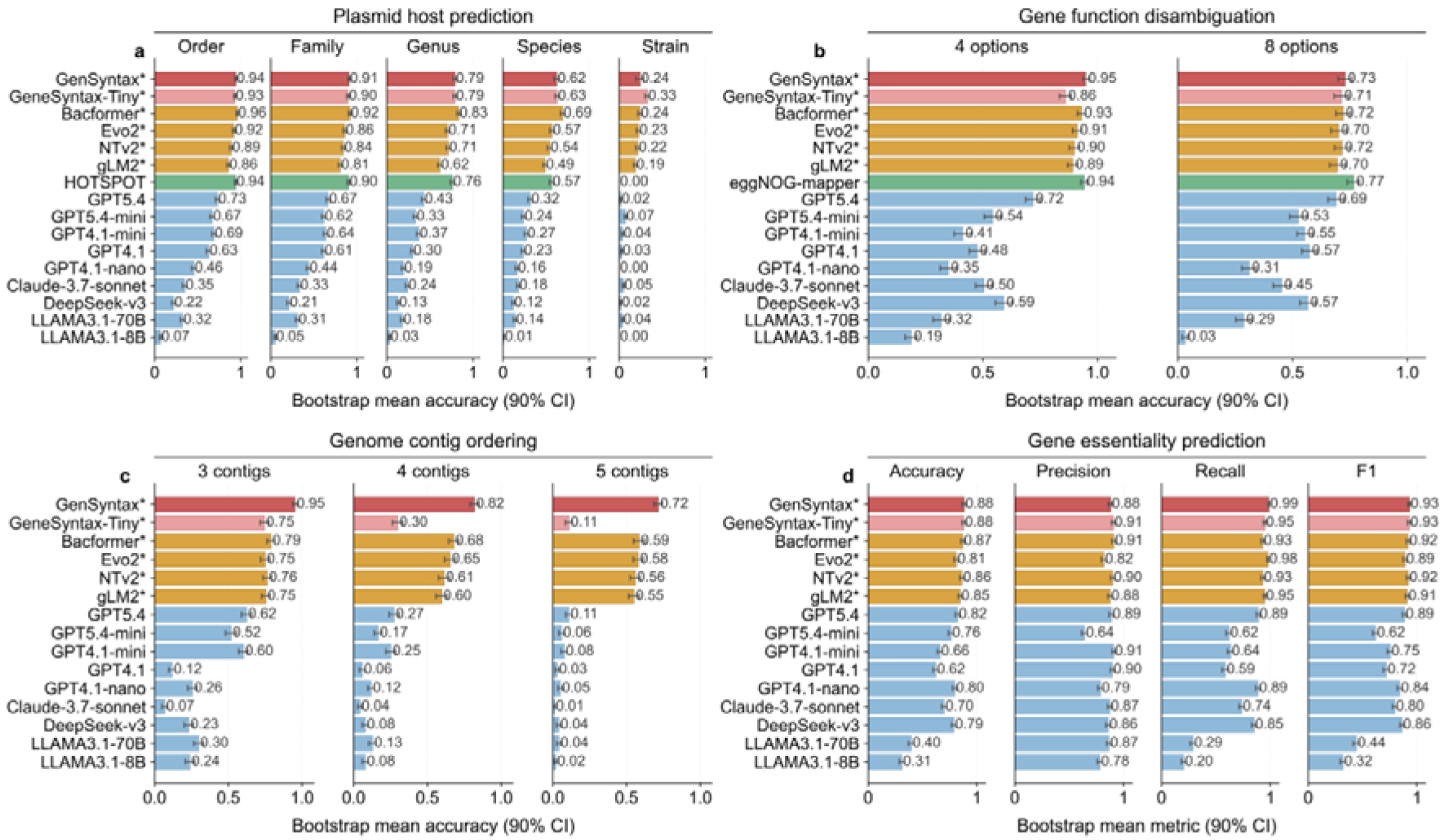
GenSyntax achieves competitive performance across four genomic prediction tasks. Performance metrics were estimated by bootstrap resampling, with points representing mean scores and error bars representing 90% confidence intervals (CIs) from 100 bootstrap replicates of the test set. Comparisons included general-purpose large language models, representative genome foundation models and domain-specific methods where applicable. **(a)** Plasmid host prediction at the order, family, genus, species and strain levels. **(b)** Gene function disambiguation from genomic product lists containing one masked gene and multiple candidate products (one correct answer and three or seven distractors). **(c)** Genome contig ordering from shuffled contig sets derived from complete chromosomal sequences. **(d)** Gene essentiality prediction from genomic context, evaluated using accuracy, precision, recall and F1 score.

To assess generalization beyond the RefSeq-derived benchmark, we further evaluated GenSyntax using an independent plasmid dataset derived from human-gut metagenomes, with host assignments inferred from CRISPR spacer matches^28,35^. GenSyntax maintained strong prediction performance across taxonomic levels, demonstrating that the learned functional representation generalizes to an independent dataset generated through a different host-assignment strategy (Supplementary Fig. S5a). We also evaluated applicability beyond complete reference genomes. Using metagenome-assembled genomes (MAGs) and incomplete draft genomes released after the GenSyntax training period, thereby eliminating potential data leakage, GenSyntax retained high prediction accuracy (Supplementary Fig. S5b and c). These results indicate that function-level genome representation learning is applicable not only to complete reference genomes but also to draft genomes and metagenome-derived assemblies.

To further assess the robustness of GenSyntax under broader annotation settings, we evaluated its sensitivity to annotation quality and annotation pipeline choice. In the pretraining corpus, hypothetical proteins accounted for 11.0% of annotated gene products, compared with 27.9% in the plasmid host prediction test set (Supplementary Fig. S6a and b). We first simulated incomplete annotations by progressively replacing annotated gene products with “*hypothetical protein”*. Prediction performance declined gradually across all taxonomic levels rather than exhibiting abrupt loss (Supplementary Fig. S6c). Even when approximately half of the input annotations were replaced, GenSyntax retained accuracies of ∼0.91, ∼0.87, and ∼0.75 at the order, family, and genus levels, respectively. Consistent with this trend, performance degradation became more pronounced at finer taxonomic resolutions, with normalized AUC loss increasing from 0.15–0.16 at the order and family levels to 0.23–0.28 at the species and strain levels (Supplementary Fig. S6d).

We further examined the effect of annotation pipeline choice by reannotating the same genomes using Prokka and Bakta^36,37^. Although PGAP-based annotations yielded the highest performance, GenSyntax retained predictive ability across alternative annotation frameworks (Supplementary Fig. S6e and f). Together, these analyses demonstrate that GenSyntax is not restricted to a single annotation pipeline, while highlighting that consistent and informative functional annotations remain important for optimal performance, particularly at fine taxonomic resolution.

Together, these results demonstrate that plasmid gene-product context contains sufficient functional information to accurately infer host identity across multiple taxonomic levels. The external validation on CRISPR spacer-linked plasmids, metagenome-assembled genomes, and incomplete draft genomes, together with the annotation robustness analyses, demonstrate that GenSyntax generalizes beyond the complete RefSeq genomes and PGAP annotations used for training, supporting its applicability to broader contexts.

### Context-specific challenge 2: Genome-contextualized gene function disambiguation

In natural language processing, word meanings are often inferable from their context, as neighboring words provide semantic clues. Similarly, in prokaryotic genomes, genomic context, *i.e.* the set of genes residing in proximity to a given gene, can provide hints about the gene’s function^19,38^. A well-known example is the operon, where functionally related genes cluster in the genome, forming coherent units analogous to “phrases” in human language^39–43^. This analogy indicates that a genome representation based on functional units may enable context-dependent inference of gene functions^19^.

To test this hypothesis, we employed a fine-tuning strategy using gene masking: one gene in the chromosome was masked, and the model was tasked with identifying the masked gene’s product within four or eight options (Fig. 1d). The challenge-specific dataset comprised 100,656 masked genes randomly selected from 50,328 chromosomes (Supplementary Fig. S2). For each masked gene, three or seven distractors were alternative products randomly sampled from the vocabulary containing more than 70,000 unique RefSeq/PGAP gene-product descriptors, while the actual functional annotations acted as the ground truth. The correct descriptor was excluded from distractor sampling, and the labels retained their original fine-grained PGAP descriptions rather than being collapsed into broad functional categories. This dataset fine-tuned the pretrained model to identify gene products from genomic context by discriminating against the distractors. We also constructed a test dataset of 1,000 masked genes randomly selected from 244 chromosomes and 256 plasmids, none of which were included in the pretraining or fine-tuning. Notably, this benchmark evaluates contextual functional disambiguation—the ability to exploit genome-wide functional context to distinguish among plausible annotations—rather than serving as a complete *de novo* annotation framework.

Under this closed-set prediction setting, GenSyntax achieved accuracies of 0.95 and 0.73 for four-option and eight-option predictions, respectively (Fig. 2b). Compared with general-purpose LLMs, genome language models, and the homology-based annotation approach eggNOG-mapper^44^, GenSyntax showed competitive performance in contextual gene-product identification. GenSyntax also retained the ability to distinguish hypothetical proteins from functionally annotated genes (Supplementary Fig. S7). Collectively, these results demonstrate that functional relationships encoded across genomic neighborhoods provide information beyond sequence similarity alone. We further evaluated GenSyntax on newly released draft genomes and MAGs that were not included during pretraining. The model retained strong performance on these datasets (Supplementary Fig. S8), supporting its applicability beyond complete reference genomes.

### Context-specific challenge 3: Genome contig ordering

As the clarity of a paragraph depends on the correct ordering of its sequences, the functional organization of a genome depends on the correct ordering of its genomic segments. We therefore asked whether GenSyntax, after learning the “grammar” of genome organization, could recover the correct order of shuffled genomic contigs based solely on their gene-product representations (Fig. 1d). We emphasize that this task addresses genome contig ordering rather than conventional *de novo* genome assembly: the inputs are predefined contigs represented by ordered gene-product descriptors, without nucleotide overlaps, sequencing reads or coverage information^45^.

To evaluate this capability, each complete chromosome was divided into three, four or five consecutive, non-overlapping contigs while preserving the internal gene order within each contig. The contigs were then randomly shuffled, and the model was trained to recover their original genomic order. Chromosome-level hold-outs were applied throughout, such that chromosomes in the test set were not exposed during pretraining (Supplementary Fig. S2). On the test set of 1,500 held-out chromosomes, GenSyntax consistently outperformed general-purpose LLMs and genome language models, achieving accuracies of 0.95, 0.82 and 0.72 for the 3-, 4- and 5-contig settings, respectively (Fig. 2c). We further evaluated GenSyntax on an independent benchmark constructed from the CAMI3 human gut long-read dataset after removing genomes overlapping with the training corpus^46^, as well as on MAGs released to RefSeq after the training period. Across both external datasets, GenSyntax maintained strong contig-ordering performance (Supplementary Fig. S9), demonstrating that the learned functional representations generalize beyond the complete RefSeq genomes used for training. Together, these results show that genome-scale functional context provides sufficient information to support accurate contig ordering and highlight the utility of function-level genome representation learning for genome organization tasks.

### Context specific challenge 4: Gene essentiality prediction

Essential genes encode gene products required for core biological processes such as metabolism, transcription, and DNA replication; their disruption can substantially impair cellular viability under defined conditions. Identifying essential genes is critical for elucidating fundamental cellular mechanisms and informing the development of targeted antimicrobial therapies^47^. To assess GenSyntax’s ability to predict gene essentiality from functional context, we curated a dataset of experimentally validated essential genes from the Database of Essential Genes (DEG)^48^, comprising over 20,000 genes from 58 bacterial and 4 archaeal species. We focused exclusively on chromosomal genes, as plasmid-encoded elements are generally considered nonessential^49^.

For each genome in DEG dataset, we retrieved the corresponding complete assembly from RefSeq and extracted the product descriptors and chromosomal coordinates for all essential genes. Each essential gene was paired with about four randomly selected nonessential gene from the same chromosome. The resulting dataset, containing both validated essential and nonessential genes, was divided into training and testing subsets at a 4:1 ratio. Each input sample included the ordered sequence sequence of gene product descriptors for the chromosome and the position of the query gene within this sequence. GenSyntax was then fine-tuned to classify genes as essential or nonessential based on their surrounding functional context (Fig. 1d).

Notably, both general-purpose LLMs and genome language models exhibited baseline ability to infer gene essentiality (Fig. 2d), suggesting an intrinsic capacity to model functional dependency. Among all evaluated models, including the newly added GPT5.4-series baselines and genome language models, GenSyntax achieved the highest recall and tied for the highest F1-score, while maintaining competitive accuracy and precision (Fig. 2d). The performance of GenSyntax remained robust under the genome-level hold-out evaluation, indicating that the model generalizes beyond genomes used during training (Supplementary Fig. S10).

To further evaluate generalization beyond the DEG dataset, we introduced external benchmarks derived from a published study on *Streptococcus suis* SC19 strain^50^: one based on experimental transposon insertion sequencing (Tn-seq) and the other on genome-scale metabolic model predictions. Neither dataset was used during training or validation. GenSyntax maintained strong performance on both external benchmarks, demonstrating that the learned functional representations generalize across independent datasets and distinct essentiality annotation strategies (Supplementary Fig. S11). Together, these results show that genome-scale functional context contains substantial information about gene essentiality and that function-level genome representation learning provides an effective framework for essentiality prediction.

### Embedding space of prokaryotic genome semantics

Following extensive training on gene product sequences, GenSyntax learned contextual patterns within prokaryotic genomes by embedding each replicon into a high-dimensional semantic space. Specifically, the model transforms each chromosome or plasmid into a 4,096-dimensional embedding vector derived from its ordered gene-product context. This embedding space captures relationships across replicons from diverse species, enabling genome-wide comparison based on learned semantics. To facilitate interpretation, we projected all replicon embeddings into two dimensions using t-SNE^51^.

In this reduced space, chromosomal embeddings occupied regions that were largely separated from plasmid embeddings, consistent with differences in their functional composition and genomic organization (Fig. 3a). A notable example is the formation of a discrete cluster comprising exclusively archaeal chromosomes (Fig. 3b), while chromosomes from phylogenetically related bacterial taxa occupy adjacent regions (Fig. 3c). These phylogenetic patterns suggest that chromosomal embeddings preserve lineage-specific functional constraints and reflect the vertically inherited genomic cores shaped by evolutionary cohesion. In contrast, plasmids exhibit a dispersed distribution across the embedding space, consistent with their status as mobile genetic elements that undergo frequent horizontal transfer across taxonomic boundaries (Fig. 3b and c).

**Figure 3.**
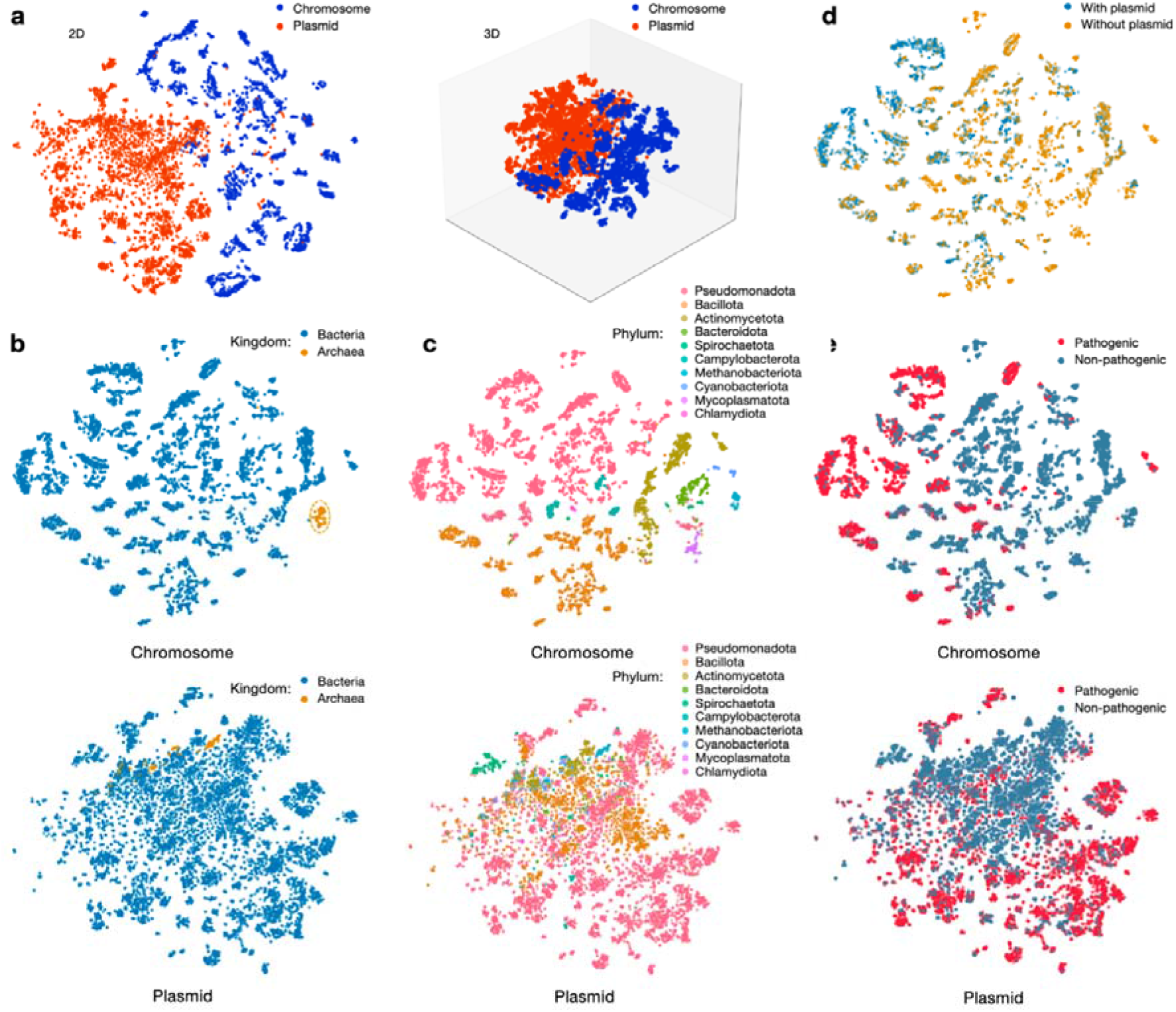
Visualization of the embedding space of genomic product sequences in GenSyntax. **(a)** 2D and 3D representations of prokaryotic replicons. Each point corresponds to a unique replicon, with distances reflecting genomic semantic similarities. Chromosomes and plasmids (shown in different colors) encoded by GenSyntax exhibit distinct spatial distribution patterns. **(b)** Archaeal chromosomes (shown in yellow) are separated from bacterial chromosomes in the embedding space. **(c)** Chromosomes from different phyla are well-separated, while plasmids exhibit overlaps. The 10 most abundant phyla in our dataset are shown in different colors. **(d)** Chromosomes of prokaryotes with plasmids and those without plasmids are visualized in different colors. **(e)** Chromosomes from pathogenic species form identifiable islands, while plasmids from pathogenic and non-pathogenic genomes show less clear spatial segregation. Pathogenic and non-pathogenic genomes are shown in blue and red, respectively.

We next examined whether GenSyntax embeddings distinguish chromosomes according to plasmid carriage. Chromosomes from genomes carrying plasmids formed partially distinct clusters from those lacking plasmids, indicating that plasmid carriage is associated with detectable differences in chromosomal functional composition (Fig. 3d). We further examined whether the model distinguishes chromosomes based on pathogenicity. Embeddings from pathogen-associated genomes showed partial enrichment in specific regions of the projected space, suggesting differences in their overall functional composition (Fig. 3e). In contrast, plasmids from pathogenic and non-pathogenic species showed overlapping distributions, consistent with the notion that plasmids carry broadly shared accessory functions, including conjugation machinery and resistance genes, which are shaped more by selective pressures than by host pathogenic status.

### Predicting microbial phenotypes from genome embeddings

The high-dimensional embeddings learned by GenSyntax encode functional semantics of prokaryotic replicons, enabling diverse downstream predictive tasks. To systematically assess the predictive utility of these representations, we compared GenSyntax with four representative genome language models (Bacformer, NT-v2, Evo2, and gLM2) on microbial phenotype prediction, including pathogenicity, plasmid carriage and experimentally validated physiological traits.

As an initial evaluation, we examined whether these genome embeddings could support accurate pathogen identification. Chromosome embeddings were generated for genomes included in the NCBI Pathogen Detection resource (Methods), and five downstream machine-learning classifiers—logistic regression (LR), random forest (RF), support vector machine (SVM), gradient boosting decision tree (GBDT) and multilayer perceptron (MLP)—were trained using identical training protocols for all embedding methods (Fig. 4a). GenSyntax consistently achieved the highest or comparable performance across all classifiers, indicating that its embeddings capture functional signatures associated with pathogenicity.

**Figure 4.**
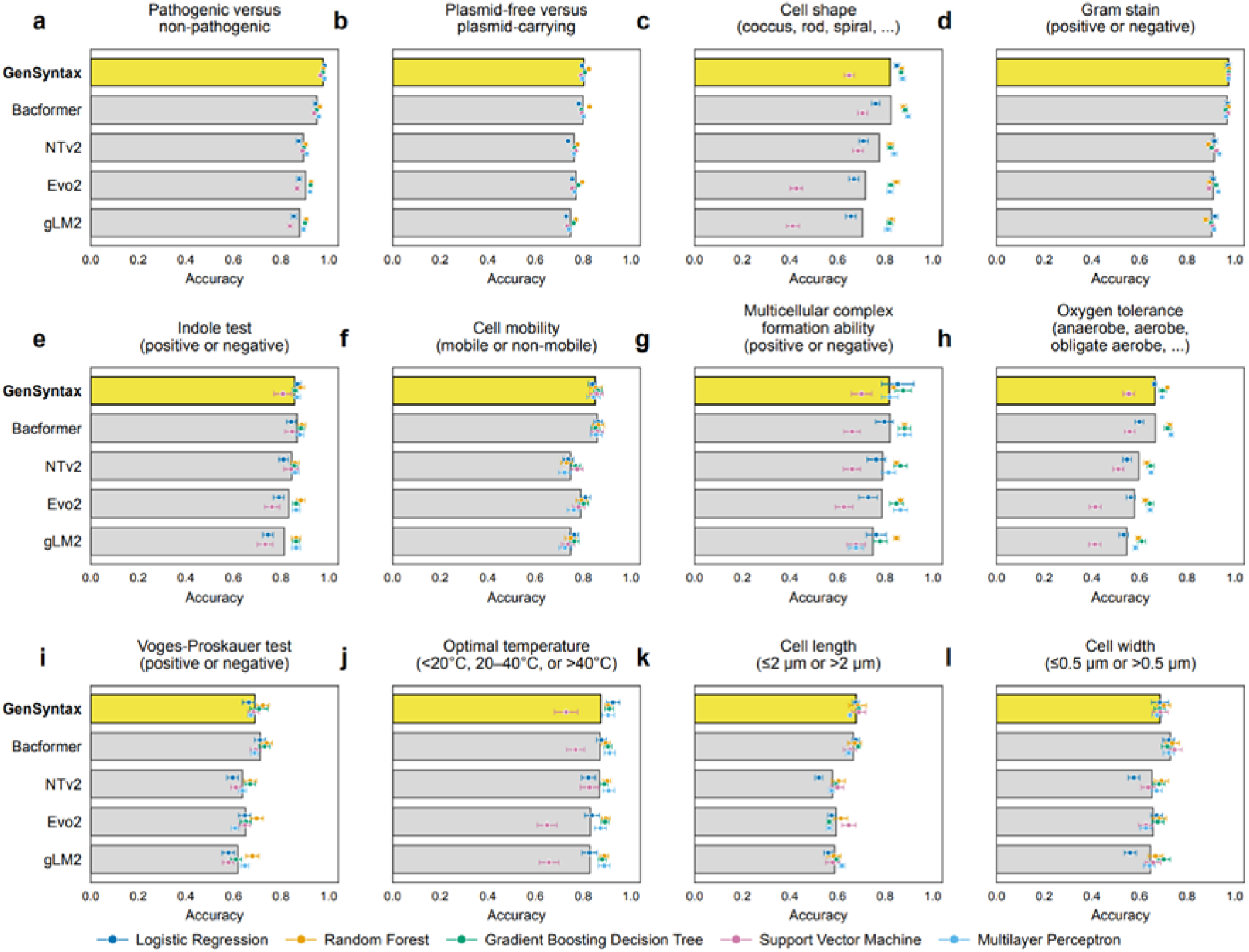
Benchmarking genome language model embeddings for microbial phenotype prediction. Chromosome-level representations generated by GenSyntax and four baseline genome language models, including Bacformer, NT-v2, Evo2, and gLM2, were used to predict a broad spectrum of microbial traits compiled from the BacDive database and other curated phenotype labels. Five machine learning classifiers were used for prediction: Logistic Regression (LR), Random Forest (RF), Support Vector Machine (SVM), Gradient Boosting Decision Tree (GBDT), and Multilayer Perceptron (MLP). Each panel shows prediction accuracies for a different phenotype or genomic trait: **(a)** distinguishing pathogenic from non-pathogenic strains, **(b)** identifying whether a genome carries plasmids, **(c)** classifying cell shape, such as coccus, rod, and spiral, **(d)** Gram staining reaction, positive versus negative, **(e)** indole test outcome, positive versus negative, **(f)** cell motility, mobile versus non-mobile, **(g)** ability to form multicellular complexes, positive versus negative, **(h)** oxygen tolerance, such as anaerobe, aerobe, and facultative anaerobe, **(i)** Voges–Proskauer test outcome, positive versus negative, **(j)** preferred growth temperature, <20 °C, 20–40 °C, or >40 °C, **(k)** average cell length, ≤2 µm versus >2 µm, and **(l)** average cell width, ≤0.5 µm versus >0.5 µm. Bar heights denote the mean accuracy across classifiers for each genome language model, while colored dots indicate individual classifier accuracies. Each dot represents the mean ± standard deviation of three replicates.

Beyond pathogenicity, we also assessed whether the chromosome embeddings contain signals related to plasmid carriage. When used to classify whether a chromosome originated from a genome with or without coexisting plasmids, the embeddings achieved classification accuracies between 0.79 and 0.82, with F1 scores in the range of 0.79 to 0.83 (Fig. 4b). GenSyntax performed competitively with Bacformer and generally outperformed NT-v2, Evo2 and gLM2, suggesting that plasmid carriage leaves detectable functional signatures within chromosomal gene-content organization.

To evaluate the broader potential of GenSyntax in phenotype inference, an important challenge in microbial genomics^52^, we expanded our analysis to a wide range of experimentally validated microbial traits. We curated a comprehensive set of phenotype annotations from the BacDive database^53^, encompassing morphology, metabolic capacity, and growth conditions. From this resource, we assembled ten phenotype-specific datasets with sufficient sample sizes (Supplementary Fig. S12), including cell shape, gram stain reaction, indole test result, motility, multicellular complex formation, oxygen tolerance, Voges–Proskauer test result, optimal growth temperature, cell length, and cell width. Species names were mapped to corresponding genome accessions to generate an integrated dataset linking genome-derived embeddings with observed phenotypes.

We then applied the five machine learning classifiers to predict phenotypes directly from GenSyntax chromosome embeddings. The model exhibited substantial predictive power across all ten traits, with greatest performances observed for Gram staining and optimal temperature (Figure 4c–4l, Supplementary Figs. S13-S15). Compared with other genome language models, GenSyntax achieved the highest or near-highest accuracy in most phenotype prediction settings, although its advantage over Bacformer was less pronounced. Together, these results demonstrate that GenSyntax embeddings capture genome-scale functional representations that extend beyond individual downstream tasks, supporting function-level genome representation learning as an effective framework for linking genomic organization with microbial phenotypes.

### Exploratory derivation of minimal genomes

Building on GenSyntax’s gene essentiality predictions, we developed an automated, context-aware pipeline for exploratory minimal genome derivation through iterative gene deletion (Fig. 5a). This approach simulates the progressive reduction of a native microbial genome by repeatedly identifying and removing nonessential genes, followed by re-evaluation of essentiality in the modified genomic context. At each step, a gene predicted to be nonessential is randomly selected for deletion from the updated pool, and this process continues until no further deletions are possible without compromising inferred viability.

**Figure 5.**
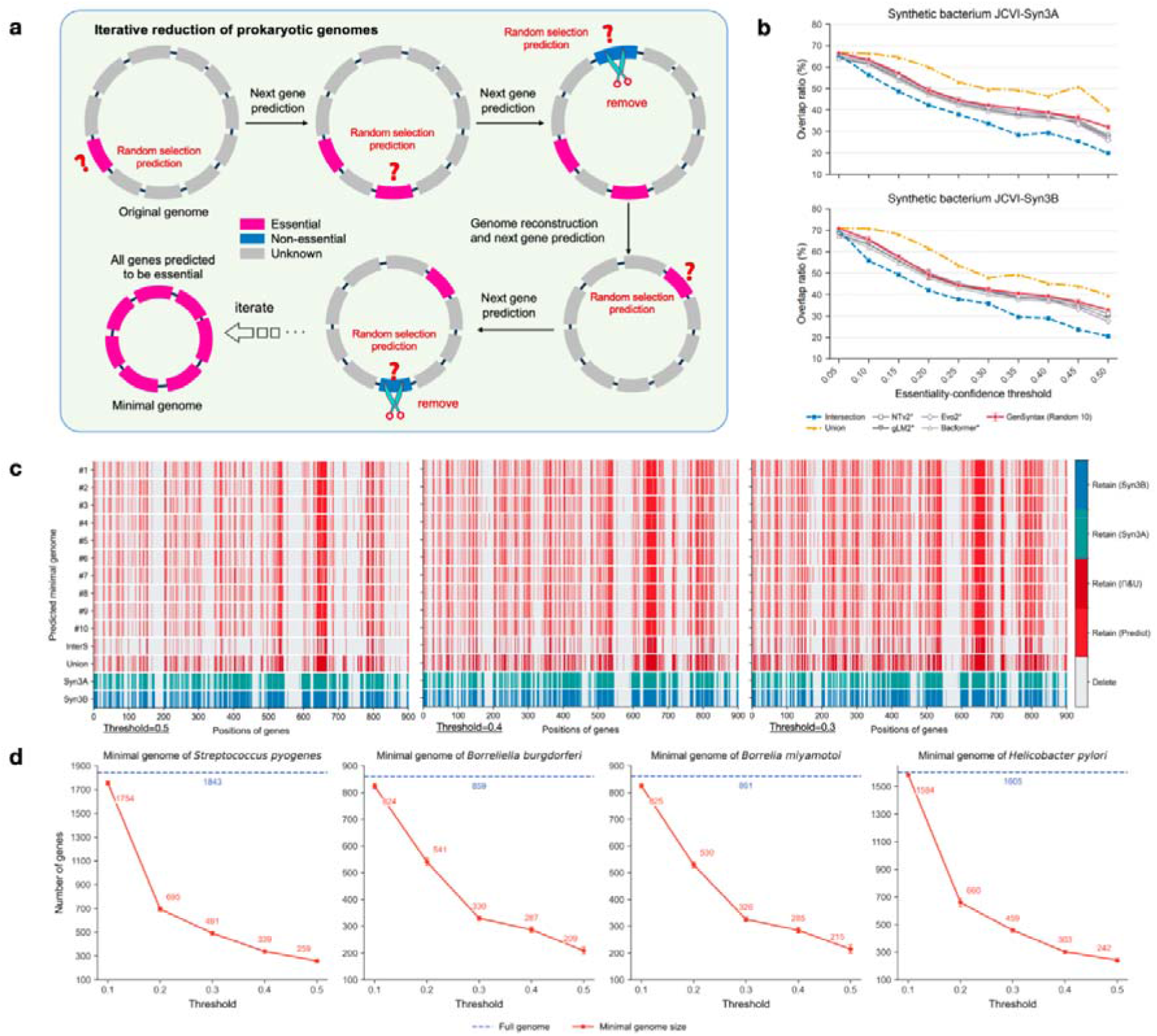
GenSyntax-enabled derivation of minimal genomes. **(a)** Schematic workflow for minimal genome derivation. **(b)** Overlap between synthetic genomes JCVI-Syn3A or JCVI-Syn3B and minimal genomes derived from genome language models. For a given genome pair, let n_1_ andn_2_ represent their respective gene counts. The number of shared genes is denoted as n*_shared_*The overlapping ratio between these two genomes is calculated 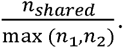 The results present the average overlapping ratio from ten randomized replicates, along with values derived from their union and intersection sets. Various confidence thresholds for gene essentiality prediction were tested. **(c)** Genomic location of retained essential genes in GenSyntax-derived minimal genomes and synthetic genomes JCVI-Syn3A and JCVI-Syn3B. Results from ten replicates, their union, and intersection were shown. Three confidence thresholds for gene essentiality prediction were tested. **(d)** Sizes of GenSyntax-derived minimal genomes across four prokaryotic species. Five confidence thresholds for gene essentiality prediction were tested for each species. Data are presented as mean standard deviation of three replicates.

We first applied this protocol to *Mycoplasma mycoides* capri GM12^54^, a well-characterized model organism for synthetic biology. Across ten independent runs of the deletion algorithm, the pipeline produced highly convergent minimal genome configurations, with greater than 90% overlap in retained genes among replicates (Fig. 5b). The resulting minimal genomes closely resembled the architecture of two experimentally validated synthetic constructs, JCVI-syn3A and JCVI-syn3B, highlighting the biological plausibility of the GenSyntax-derived designs^55^ (Fig. 5b and c).

The size and composition of the resulting minimal genomes were strongly influenced by the essentiality confidence threshold (Figure 5b and 5c). At higher thresholds, the essential-gene assignment became more stringent. Thus, more aggressive deletion was permitted, producing genome sizes substantially smaller than JCVI-syn3A/3B. In contrast, lower thresholds led to more conservative designs, retaining a broader set of genes to account for potential context-dependent essentiality. Systematic comparison against the JCVI constructs showed that lower thresholds yielded greater overlap, with 70% of retained genes matching JCVI-syn3A/3B at a threshold of 0.05 (Fig. 5b). To place this result into a broader context, we applied the same iterative reduction procedure, prediction thresholds, and evaluation criteria to NT-v2, gLM2, Evo2, and Bacformer. Across multiple confidence thresholds, GenSyntax consistently achieved larger or comparable overlap with the experimentally derived JCVI genomes (Fig. 5c), demonstrating that its function-level genome representations effectively recover known minimal genome components.

To further examine whether the derived minimal gene sets were biologically meaningful, we compared the functional composition of retained and excluded genes between the inferred gene sets and the JCVI genomes (Supplementary Table S1)^44^. Core cellular processes, including translation, protein export, DNA replication, and DNA topology maintenance, were consistently retained across both GenSyntax-derived and JCVI reduced genomes. In contrast, genes differing between the two sets were primarily associated with metabolism, membrane transport, lipoproteins, mobile genetic elements, and proteins of unknown function, categories that are often influenced by environmental conditions, functional redundancy, or annotation uncertainty. These analyses support the biological consistency and plausibility of the inferred gene sets while highlighting the context-dependent nature of genome reduction.

To evaluate generalizability, we applied the same pipeline to four additional bacterial species from distinct genera: *Borrelia burgdorferi*, *Streptococcus pyogenes*, *Borrelia miyamotoi*, and *Helicobacter pylori*. Across all species, independent runs consistently converged on similar candidate reduced gene sets, with <3% variation in retained genes (Fig. 5d), indicating that the iterative deletion procedure was reproducible under the tested settings. Strikingly, under the most aggressive threshold (threshold = 0.5), the resulting genome sizes converged within a narrow range (209-259 genes), despite wide phylogenetic divergence. This convergence suggests the existence of a conserved functional core across diverse prokaryotic lineages, while also demonstrating the pipeline’s ability to incorporate lineage-specific constraints through adjustable confidence parameters.

## Discussion

Here, we present GenSyntax, a framework for function-level genome representation learning that models prokaryotic genomes as ordered sequences of gene product descriptors rather than directly modeling nucleotide or protein sequences. By representing annotated replicons as “genetic paragraphs”, GenSyntax provides a semantic abstraction of genome organization in which each unit corresponds to a biologically interpretable functional concept. This abstraction substantially compresses genomic information while preserving functional context, enabling genome-scale modeling of genome organization and functional relationships.

Existing genome language models represent biological information at different levels of abstraction, including nucleotide sequences, protein sequences, learned embeddings or structured biological features. Models such as Evo, Evo2, and gLM2 have advanced sequence-based genome representation learning^7,13,14,17,18^, while related approaches including PlasGO, gLM and Bacformer have explored protein-level representations and genomic functional modeling^15–17^. Miller *et al.* introduced NLP-inspired representations, providing another perspective on genomic abstraction^19^. Recent advances, particularly Evo2, have further demonstrated the feasibility of long-context nucleotide modeling at genome scale^21^. In this landscape, GenSyntax introduces a distinct representation strategy by using human-readable gene product descriptors as the fundamental modeling units. Rather than learning genome organization primarily from sequence patterns, GenSyntax captures genome structure through functional semantics, providing a compressed and interpretable representation that directly connects model outputs with biological annotations. Thus, the contribution of GenSyntax is not to replace existing sequence-based genome language models, but to establish a complementary functional representation paradigm that enables genome-scale reasoning through biologically meaningful semantic units.

Compared with many existing genomic foundation models trained on substantially larger collections of nucleotide or protein sequences, GenSyntax was adapted using a relatively compact corpus of 49,250 annotated prokaryotic genomes (Supplementary Fig. S16). Despite this difference in genomic training scale, GenSyntax achieved competitive performance across multiple genome-scale tasks. These results suggests that representation choice can substantially influence the efficiency of genomic foundation model training: by operating on gene product descriptors that directly encode functional semantics, GenSyntax focuses model capacity on genome-level functional organization rather than requiring the model to infer biological meaning solely from large-scale sequence patterns. Although training data scale and representation modality are not directly comparable across models, these results highlight the potential of semantic genome representations as a data-efficient strategy for functional genome modeling.

The minimal genome analysis illustrates one potential downstream application of this representation. Using an iterative context-aware gene deletion strategy, GenSyntax generated highly reproducible reduced genome designs that showed substantial overlap with experimentally constructed JCVI synthetic genomes. Functional analyses further showed that conserved genes were enriched for core cellular processes, whereas most differences involved condition-dependent metabolic functions, membrane-associated proteins, mobile genetic elements, and genes with unknown functions. These findings support the biological plausibility of the predicted reduced genomes, although we emphasize that they should be regarded as hypothesis-generating computational predictions rather than experimentally validated genome designs^56^.

By centering on human-interpretable semantics, GenSyntax improves the readability of genome-scale model inputs and outputs compared with raw nucleotide or protein sequences. Its product-level representation provides an intuitive framework for integrating genomic data with biological knowledge, fostering accessibility and collaboration across disciplines. To further improve practical usability, we developed a web interface (http://111.2.199.31:2103/) that supports single-sample and batch prediction across the main GenSyntax tasks. We also introduced GenSyntax-Tiny, a lightweight 0.6B-parameter model based on Qwen3, which reduces computational requirements while retaining competitive performance across multiple benchmarks. These developments improve practical accessibility of GenSyntax.

An important characteristic of GenSyntax is that it operates on functional annotations rather than raw nucleotide sequences. Consequently, the framework is most applicable to well-annotated prokaryotic genomes, plasmids, draft genomes, metagenome-assembled genomes, and functionally annotated contigs. It is not intended for raw sequencing reads, unannotated assemblies, or applications requiring nucleotide-level resolution, such as variant-effect prediction or motif discovery. Our robustness analyses further indicate that GenSyntax remains effective across different annotation pipelines and under moderate annotation degradation, although performance decreases as annotation quality declines. These observations define both the strengths and the current boundaries of annotation-driven genome modeling.

Several limitations remain. First, the current model was developed using an 8-billion-parameter foundation model, and larger architectures or larger training dataset may further improve the modeling of complex functional relationships. Second, given the extensive evolutionary relatedness among microbial genomes, validation using biologically independent datasets provides the most stringent assessment of whether GenSyntax learns transferable functional representations. Although current external validation substantially broadens the evaluation, additional benchmarking across more diverse environmental microbiomes and taxonomic groups will further establish generalizability. Third, GenSyntax models the information encoded in current functional annotation systems; improvements in annotation quality will directly influence future versions. Forth, because GenSyntax models genomes exclusively through functional annotations, it cannot capture nucleotide-level regulatory information or sequence variation that is absent from gene-product descriptions.

To address these limitations, future work may advance along three complementary directions. First, scaling GenSyntax to larger foundation models could enhance its capacity to model complex and subtle genomic features. Second, further expanding the training corpus to include partially annotated genomes, metagenomes, and even eukaryotic genomes would improve generalizability and extend applicability beyond prokaryotes. Third, integrating semantic representations of GenSyntax with nucleotide-based genomic foundation models may provide a unified framework that combines the interpretability and efficiency of functional annotations with the molecular resolution of sequence-based models. Such multimodal genome foundation models may enable more comprehensive genome-scale reasoning across multiple levels of biological organization.

## Methods

### Curation of prokaryotic genome dataset

We curated a dataset of 49,250 fully sequenced prokaryotic genomes from the NCBI RefSeq database as of April 18, 2025^24^, limiting the selection to assemblies explicitly labeled as “complete”. All genomes were annotated using the Prokaryotic Genome Annotation Pipeline (PGAP), which identifies protein-coding sequences (CDSs) and various noncoding RNA elements, including ncRNAs, tRNAs, rRNAs, and tmRNAs^22^. Detailed gene annotations were obtained from the corresponding GBFF (GenBank Flat File) files, which include functional descriptions, genomic coordinates, and replicon-level metadata (e.g., classification as “chromosome” or “plasmid”), along with associated host taxonomy. Using these GBFF files, we developed custom scripts to systematically extract all annotated gene product names from each replicon and to retrieve host organism information for each plasmid.

### Continuous pre-training

The foundation LLM was initially trained on a broad corpus of human-curated textual data, encompassing both general and domain-specific knowledge, including genomic content. To further adapt the model to the contextual semantics and syntax of gene product sequences, we constructed a genome-ordered corpus of gene product descriptors. The primary GenSyntax model was initialized from LLaMA3.1-8B and further adapted using this gene-product corpus. This corpus included 50,328 chromosomal paragraphs (mean length: 3,754 gene products; maximum: 12,898) and 59,613 plasmid paragraphs (mean length: 108; maximum: 5,201). Plasmids with fewer than three annotated products (n = 145) were excluded due to insufficient sequence length for effective training. All paragraphs were tokenized using the LLaMA 3.1 tokenizer and used for continuous pre-training.

In the pretraining corpus, hypothetical proteins accounted for 10.75% of annotated gene products. These genes were retained without filtering or imputation and were represented using their original annotation text, just like all other gene products. Consequently, multiple hypothetical proteins shared the same textual token. Rather than relying exclusively on the identity of individual product annotations, GenSyntax learns contextual representations from the ordered co-occurrence of neighboring genes across complete genomes.

Training was performed across five servers, each equipped with eight H800 GPUs. The key hyperparameters were: sequence length of 128,000 tokens, learning rate of 1.0 × 10, per-device batch size of 1, gradient accumulation steps of 4, and one training epoch. To support efficient long-context training, we utilized parameter-efficient fine-tuning via Low-Rank Adaptation (LoRA) and applied flash attention mechanisms to reduce memory overhead and accelerate convergence. No large-scale hyperparameter optimization was performed; instead, training configurations were selected based on the recommended settings of the base models and training framework, with limited adjustments to ensure stable and reproducible optimization.

In addition to the full-scale GenSyntax model, we developed GenSyntax-Tiny to evaluate whether the proposed gene-product representation remained effective independent of large model capacity. GenSyntax-Tiny was initialized from Qwen3-0.6B and trained using the same gene-product-based genomic adaptation strategy. Both GenSyntax and GenSyntax-Tiny were subsequently evaluated on task-specific datasets through downstream fine-tuning following predefined training protocols.

### Construction of the plasmid host prediction dataset

To construct the plasmid host prediction benchmark, we curated all prokaryotic plasmids with assigned host taxonomic labels from RefSeq genomes. To ensure sufficient genomic context for functional representation learning, plasmids were retained only if they contained at least four annotated genes. Host labels were collected at multiple taxonomic resolutions, including order, family, genus, species, and strain levels, enabling evaluation of GenSyntax across hierarchical host classification tasks.

Among the curated plasmids, 59,613 plasmids were used for task-specific fine-tuning, allowing GenSyntax to learn associations between plasmid functional composition and host identity across different taxonomic levels. The remaining 1,000 plasmids were reserved as test set and were not used during fine-tuning. Importantly, all plasmids in the test set were completely excluded from the genomic pretraining corpus, ensuring that evaluation was performed on previously unseen plasmid sequences.

### Curation of an IMG/PR human gut metagenomic plasmid benchmark

To evaluate plasmid host prediction on an independent benchmark with host assignments, we curated plasmids from the IMG/PR dataset^28,35^. Plasmid records were retained if they satisfied all of the following criteria: (i) classified as complete plasmid sequences; (ii) recovered from metagenomic samples; (iii) originating from human gut microbiome fecal samples; and (iv) having CRISPR spacer-based host assignments available at the species level. After quality filtering, 1,314 complete plasmids with species-level host assignments were retained for evaluation. All plasmids were independently reannotated using Bakka, and the resulting gene-product annotations were used as GenSyntax input. Host prediction was performed directly using the pretrained model without additional fine-tuning, model adaptation, or parameter optimization on this dataset.

### Construction of gene-product disambiguation dataset

In the 4-option and 8-option gene function disambiguation tasks, the correct option corresponds to the original gene product descriptor of the masked gene annotated by RefSeq/PGAP. The remaining distractor options were randomly sampled from the complete gene product vocabulary extracted from NCBI RefSeq-annotated genomes, which contains more than 70,000 unique gene product descriptors. The correct label was excluded during sampling to ensure that each candidate set contained only one valid answer. Regarding label granularity, we retained the original PGAP gene product annotations without manually collapsing them into broader functional categories. Therefore, the prediction targets generally represent fine-grained functional descriptions, such as “ferric enterobactin transporter” or “ABC transporter permease,” rather than broad functional classes such as “transporter”.

### Construction of genome-contig ordering dataset

For the task of genome contig ordering, each complete chromosome was represented as an ordered sequence of gene-product descriptors according to genomic coordinates. The chromosome sequence was then partitioned into 3, 4, or 5 non-overlapping contigs, with the internal order of genes within each contig preserved. For the 3-, 4-, and 5-contig settings, contigs contained means of 671.52, 503.64, and 402.91 gene products, with size ranges of 3–4300, 2–3225, and 2–2580, respectively. The resulting contigs were randomly shuffled before being provided as model input, and the task was defined as recovering the original genomic order. Because this task evaluates genome organization from functional context rather than sequence-overlap information, no artificial overlap was introduced between contigs. For circular chromosomes, we considered cyclically equivalent contig arrangements as correct to avoid penalizing predictions that differed only due to the arbitrary choice of chromosome starting position.

### Curation of gene essentiality dataset

To assess GenSyntax’s ability to predict gene essentiality, we curated a dataset of over 20,000 experimentally validated essential genes from the Database of Essential Genes, covering 58 bacterial and 4 archaeal species^48^. Only chromosomal genes were included, as plasmids are typically considered nonessential accessory elements. For each genome listed in the database, we retrieved the corresponding complete RefSeq assembly using its accession number. Essential genes were then mapped to their reference assemblies based on chromosomal coordinates, and their associated gene product descriptors were manually extracted for downstream analysis.

### Supervised fine-tuning

Following continuous pre-training, the model had gained an initial grasp of the semantics and syntax of prokaryotic genomes but lacks the ability to perform specific genomics tasks. To address this, we designed four supervised fine-tuning tasks using the curated dataset: (1) predicting plasmid host from product profiles (59,613 plasmids); (2) predicting masked gene products from replicon product lists (50,328 chromosomes, 100,656 masked genes); (3) context-aware ordering of genomic assembly sequence (1,080,000 contigs from 50,328 chromosomes); and (4) predicting gene essentiality from learned genomic language patterns (509 chromosomes, 23,532 essential genes, and 91,514 non-essential genes). Training was conducted under the same environment and hyperparameters as in the pre-training phase, except that the loss function was computed only on the model outputs. We trained for three epochs, with data from Task 1 duplicated in the training set to compensate for its relatively smaller size.

For the task of predicting masked gene products, GenSyntax outputs the most probable gene product when provided with the corresponding instructions and inputs. For the plasmid host prediction task, the model outputs the predicted host in the format [order, family, genus, species, strain]. For the genomic assembly ordering task, GenSyntax returns the predicted sequence of contigs as [Contig X–Contig Y– … –Contig Z]. For the gene essentiality prediction task, the model outputs whether the queried gene is classified as [essential] or [non-essential].

### Implementation and task-specific training of genomic language model baselines

We evaluated four pretrained genomic language model baselines: nucleotide-transformer-v2-500m-multi-species (NT-v2-500M), gLM2-650M, Evo 2-7B and Bacformer-26M. Each model was loaded from its publicly released checkpoint using the tokenizer, preprocessing pipeline and basic inference settings provided by its official repository. NT-v2, gLM2 and Evo 2 received nucleotide-sequence inputs, whereas Bacformer received the corresponding amino-acid sequences. For all downstream tasks, the pretrained model parameters were frozen and each model was used exclusively as a feature encoder. The extracted representations were supplied to task-specific multilayer perceptron (MLP) prediction heads, with gradients propagated only through the MLP parameters. The same biological samples, labels and train–test partitions were used across models.

For NT-v2, sequences were processed using its 6-mer tokenizer, and the final hidden-layer representations were mean-pooled over valid non-padding positions. For gLM2, nucleotide sequences were processed with its released tokenizer, and last_hidden_state was similarly mean-pooled. For Evo 2, nucleotide sequences were encoded at single-nucleotide resolution, and representations were extracted from the intermediate layer blocks.28.mlp.l3, following the embedding procedure recommended in its official repository, before mean pooling. For Bacformer, each protein was first embedded using ESM-2 (esm2_t12_35M_UR50D) by averaging its amino-acid token representations. The ordered protein embeddings were subsequently processed by Bacformer, and the final contextual protein representations were mean-pooled over valid protein positions. Inputs exceeding a model’s maximum supported length were truncated to the corresponding model-specific limit, and padding positions were excluded from pooling.

For plasmid host identification, five independent multiclass MLP classifiers were trained for the order, family, genus, species and strain levels. For gene-product disambiguation, each target gene was associated with one correct candidate and either three or seven randomly sampled distractors. Each candidate was paired with the same genomic context and classified as matching or non-matching. Separate binary classifiers were trained for the four-option and eight-option settings. During inference, all candidates were scored independently, and the candidate with the highest positive-class probability was selected as the predicted gene product.

For genome contig ordering, separate multiclass classifiers were trained for the three-, four- and five-contig settings. For n input contigs, the label space contained all n! possible permutations. Because chromosomes were treated as circular, cyclic rotations of the reference order were considered equivalent during evaluation. For gene essentiality prediction, binary classifiers were trained using the target-gene nucleotide representations from NT-v2, gLM2 or Evo 2, or the corresponding amino-acid representations from Bacformer.

### Output Formatting and Label Parsing

For all tasks, prompts explicitly required GenSyntax to return only the final answer in a predefined format. After fine-tuning, the model produced valid standard-format outputs in most cases; for gene essentiality prediction, more than 99% of outputs directly matched the allowed labels {essential, non-essential}.

Model outputs were parsed using deterministic rules fixed before evaluation, with no manual correction. For plasmid host identification, predicted host names were normalized by lowercasing and removing non-alphabetic characters, and then matched at the corresponding taxonomic rank. For gene product disambiguation, option labels were extracted using regular expressions, prioritizing bracketed answers such as [A], followed by numbered answers, explicit “Answer: A” patterns and standalone option letters. For genome contig ordering, contig identifiers were extracted by regular expression, and cyclically equivalent orders were treated as correct for circular genomes. For gene essentiality prediction, outputs were normalized to lowercase and mapped to “essential” or “non-essential”, with “non-essential” matched first to avoid substring errors.

Outputs that could not be mapped to a valid label were marked as invalid and included in the evaluation statistics. The same parsing rules were applied consistently across models, ensuring reproducible conversion from natural-language outputs to discrete evaluation labels.

### Hypothetical protein recognition

To evaluate whether GenSyntax can identify genes lacking sufficient functional characterization, we constructed a masked product recognition task using genes with either functional product annotations or “hypothetical protein” annotations. The original gene product descriptors were randomly masked, and the model was instructed to predict whether the masked product corresponded to a hypothetical protein or a functionally annotated gene. The task was evaluated as a binary classification problem, with predictions mapped to the labels “hypothetical protein” and “non-hypothetical protein” using predefined output parsing rules. This evaluation tests whether GenSyntax can leverage genome-wide functional context to distinguish genes with limited annotation evidence from those with established functional descriptions, thereby assessing its ability to recognize annotation uncertainty.

### Assessment of robustness to annotation quality

We evaluated the robustness of GenSyntax to annotation quality in the plasmid host prediction task using two perturbation schemes. First, to simulate annotation degradation, we progressively increased the proportion of hypothetical protein annotations in the test set from the original 28% to 98% by randomly replacing annotated gene product descriptors while preserving genome identities and taxonomic labels. Prediction accuracy was evaluated at the order, family, genus, species, and strain levels for each perturbation level. Overall sensitivity to annotation degradation was quantified as the normalized area-under-the-curve (AUC) loss relative to the unperturbed test set.

Second, to evaluate robustness to annotation pipeline differences, we reannotated the same plasmid test set using Prokka, and Bakta^36,37^. Because both continuous pre-training and supervised fine-tuning were performed using PGAP annotations, PGAP served as the reference annotation. The resulting gene-product sequences were evaluated under identical inference settings, and prediction accuracies across the five taxonomic ranks were compared among annotation pipelines. Together, these analyses evaluated the robustness of GenSyntax to both incomplete functional annotation and variation in annotation pipelines.

### Curation of post-training draft genomes and metagenome-assembled genomes

To evaluate temporal and cross-dataset generalization, we curated two independent collections of prokaryotic assemblies released after the GenSyntax pre-training cutoff (18 April 2025). The first collection comprised 500 RefSeq draft genomes released in 2026 with estimated completeness between 80% and 100%. The second collection comprised 142 metagenome-assembled genomes (MAGs) released after the cutoff date and containing RefSeq annotations. All assemblies were downloaded from NCBI in GenBank flat-file (GBFF) format, and their annotated CDS, tRNA, rRNA, ncRNA and tmRNA features were extracted in genomic order.

For plasmid host identification, we retained only replicons explicitly annotated as plasmids and containing at least four annotated gene products, resulting in 49 plasmids from the draft-genome collection and 73 plasmids from the MAG collection. The ordered gene-product descriptors of each plasmid were used as input, and the source organism taxa was used as the host label. Host prediction was evaluated at taxonomic ranks from class to species because reliable strain-level annotations were unavailable for most records.

For gene function prediction, non-plasmid contigs within each assembly were concatenated to preserve genomic context. Assemblies containing fewer than 20 annotated gene products were excluded. One CDS per assembly was randomly masked to generate evaluation instances, yielding 200 masked-gene instances from draft genomes and 150 from MAGs. Four-choice and eight-choice prediction tasks were constructed for each masked gene, with distractor candidates randomly sampled from the total product vocabulary. Neither dataset was used for pre-training or task-specific fine-tuning; both were reserved exclusively for post-training external evaluation.

For contig ordering evaluation, the MAG collection was additionally used to assess the generalizability of GenSyntax for recovering genomic organization from fragmented assemblies. Each MAG was divided into 3, 4, or 5 non-overlapping contigs while preserving the original genomic coordinates, followed by random permutation of contig order.

### Gene function prediction based on eggNOG-mapper

As a conventional sequence- and orthology-based baseline, the amino acid sequence of each target protein was annotated using eggNOG-mapper in DIAMOND mode^44^. The resulting ‘Description’ and ‘Preferred_name’ fields were extracted and converted into the same four- or eight-choice prediction format used for GenSyntax through a deterministic, training-free text-matching procedure.

Briefly, eggNOG-mapper annotations and candidate gene-product names were normalized by lowercasing, replacing punctuation and hyphens with spaces, collapsing redundant whitespace, and removing non-specific qualifiers (for example, *putative*, *probable*, *possible*, and *predicted*). Similarity was then calculated independently at the word and character levels using Sørensen-Dice coefficients over word sets and character trigrams^57^, respectively, and the two scores were equally weighted. Each candidate was compared separately with the eggNOG-mapper functional description and preferred gene name, and the higher of the two similarity scores was used as its final matching score. The candidate with the highest score was selected as the prediction.

Queries lacking valid eggNOG-mapper annotations or yielding zero similarity with all candidates were classified as unresolved and counted as incorrect in accuracy calculations. All preprocessing rules, similarity metrics, and decision criteria were predefined before evaluation and were not optimized using ground-truth labels.

### External benchmarks for genome contig ordering and gene essentiality prediction

An independent genome contig ordering benchmark was constructed from the CAMI3 human gut toy long-read dataset (sample 19 GSA; https://s3.bi.denbi.de/swift/v1/cami/cami3_toydata/human-gut-toy-long/sample_19_gsa.tar.gz)^46^. Genome files in this dataset were first parsed to extract NCBI accession identifiers from the FASTA headers, yielding 407 genomes with valid accession information. The corresponding RefSeq GBFF annotation files were downloaded from NCBI and parsed to obtain ordered gene-product annotations, nucleotide sequences and genomic coordinates. To avoid overlap with the original training corpus, genomes already present in the training set were removed, leaving 322 non-redundant genomes. Genomes containing fewer than 20 annotated gene products were then excluded, resulting in 287 genomes for the final CAMI-287 benchmark. For each retained genome, ordered gene-product sequences were partitioned into three, four or five contiguous blocks, which were randomly shuffled to form the input contig sets. The original genomic order was used as the ground-truth label, and cyclically equivalent predictions were treated as correct.

Two independent gene essentiality benchmarks were derived from a published *Streptococcus suis* study integrating transposon insertion sequencing (Tn-seq) and genome-scale metabolic modeling^50^. Essential and non-essential gene labels were obtained separately from metabolic-model predictions and experimental Tn-seq screening. Gene identifiers from both datasets were mapped to the corresponding RefSeq annotation, and genes with ambiguous labels or unreliable mappings were excluded.

To enable comparison with genomic language models that operate on different molecular representations, each benchmark was formatted in parallel as gene-product, nucleotide-sequence and amino-acid-sequence versions whenever applicable. The gene-product version was used for GenSyntax, whereas nucleotide- or protein-based versions were used for corresponding genomic language model baselines according to their expected input modality. The curated CAMI-287 and gene-essentiality datasets were not used during model training or validation and were used exclusively as independent external benchmarks to assess cross-dataset generalization.

### Compilation of pathogenic genome list

To map pathogenic prokaryotic genomes within the embedding space, we utilized the NCBI Pathogen Detection resource, which provides a global repository of pathogenic genome sequences. As of August 19, 2025, we retrieved the full genome list and applied stringent filters to retain only genomes that met the following criteria: (1) labeled as “complete” and (2) derived from “clinical” isolates. This filtering process yielded 29,021 high-confidence pathogenic entries. Each genome was then matched to its corresponding RefSeq record using the provided assembly accession numbers.

### Predicting microbial phenotypes from genome embeddings

To evaluate the capacity of GenSyntax embeddings to predict phenotypic traits, we retrieved ten microbial phenotype datasets from the BacDive database using the Advanced Search interface. We prioritized traits with sufficient sample sizes to support robust modeling. In total, we assembled datasets for cell shape (n = 8,282), gram staining result (n = 8,459), indole test outcome (n = 3,419), cell mobility (n = 8,126), multicellular complex formation (n = 1,561), oxygen tolerance (n = 15,262), Voges-Proskauer test results (n = 3,316), optimal temperature (n = 5,557), cell length (n = 4,376), cell width (n = 4,253). Species names were matched to genome accessions in RefSeq to construct integrated genotype–phenotype datasets, yielding 2,352, 2,464, 1,231, 2,371, 291, 2,837, 1,090, 1,291, 1,034, and 987 usable entries for the ten phenotypes, respectively.

These curated datasets encompass categorical, multiclass, and numerical phenotypes. Of the ten traits, four were binary (cell motility, multicellular complex formation, indole test outcome, and Voges–Proskauer test outcome). Gram staining results comprised three classes (positive, negative, variable), while oxygen tolerance included nine categories (anaerobe, facultative anaerobe, obligate anaerobe, microaerophile, aerobe, obligate aerobe, facultative aerobe, microaerotolerant, or aerotolerant). Cell shape comprised twenty categories (coccus, rod, filament, oval, ellipsoidal, ovoid, curved, sphere, pleomorphic, spiral, vibrio, helical, flask, dumbbell, ring, spore, crescent, star, diplococcus, or other). Three traits—optimal temperature, cell length, and cell width—were recorded as numerical values. For traits expressed as ranges (e.g., cell length 1–2.5 μm), the midpoint was calculated. The continuous variables were discretized into biologically meaningful classes: optimal temperature (<20°C, 20–40°C or >40°C), cell length (≤2 μm or >2 μm), and cell width (≤0.5 μm or >0.5 μm).

To predict microbial phenotypes, we trained five machine learning models, logistic regression, random forest, support vector machine, gradient boosting decision tree, and multilayer perceptron, using genome embeddings as input features. Each genotype-phenotype dataset was randomly split into training and test sets in a 4:1 ratio. Model performance was evaluated using both accuracy and weighted F1 score to account for class imbalance.

### Derivation of minimal genomes

To systematically derive the minimal genome, we propose an iterative reduction algorithm (IRA) that leverages GenSyntax’s ability to predict gene essentiality. By progressively eliminating genes predicted to be non-essential until no further reduction is possible, the algorithm ultimately obtains a minimal yet viable genomic structure. Let the complete genome be represented by a set of genes (G= {g_1_, g_2_,…, g_3_}), with corresponding product sequences (*P*^(0)^ = {P_1_, P_2,_ …, P_n_}) where P_i_ is the encoded product of g_i_. The gene essentiality prediction function of the GenSyntax model is defined as: GenSyntax, P 0,1, where 1 indicates that the gene g_i_ corresponding to product P_i_ is essential in the genomic context represented by the current product set P; 0 indicates that the gene is non-essential and can be removed. Since the model can only process one product at a time, the IRA algorithm employs a random selection mechanism to enable iterative reduction.

The algorithm proceeds in discrete iterations indexed by t, with the following steps:

1. Initialization: Start with the complete product set *P*^(0)^, corresponding to the wild-type genome.
2. At iteration t, randomly select a product P_i_from the remaining product set *P*^(t)^, and evaluate the essentiality of its corresponding gene using GenSyntax;
3. If the output is 0 (non-essential), remove the product and its corresponding gene, update the product set to*P*^(t+1)^ = *P*^(t)^\ (P_i_) reset t ←t+1, and proceed to the next iteration.
4. If the output is 1 (essential), return to the remaining product set, randomly select another product, and repeat the evaluation until a removable non-essential gene is identified.

The IRA algorithm terminates when a random traversal evaluation of all gene products in the remaining set *P*^(t)^ shows that all their corresponding genes are essential (i.e.,*P*_i_∈ *P*_min_) *P*^(t)^, GenSyntax (*P*^(t)^, P_i_) = 1)) At this point, the final set *P*_min_ *P*^(t)^ corresponds to the minimal genome G_min_ = {g_i_ | P_i_ ∈ *P*_min_} This algorithm adapts to GenSyntax’s constraint of processing only one product per inference through a random selection mechanism, ensuring local minimality under the model’s predictions—meaning no single gene can be removed to obtain a more compact viable genome. The random traversal strategy avoids biases from evaluation order and efficiently converges to the minimal solution within the vast combinatorial space of possible genomes.

## Data Availability

All data associated with this work are available at the Github repository (https://github.com/nishiwen1214/GenSyntax).

## Code Availability

All codes associated with this work are available at the Github repository (https://github.com/nishiwen1214/GenSyntax).

### Acknowledgement

This study was supported by the National Key R&D Program of China (2024YFA0920200 to TW), the National Natural Science Foundation of China (32470701 and 12401660 to TW), and the Shenzhen Institute of Synthetic Biology Scientific Research Program (HSE499011086 to TW). Shiwen Ni was supported by Shenzhen Science and Technology Program (JCYJ20250604182917023), Shenzhen Major Science and Technology Project (KCXFZ20240903094007010), GuangDong Basic and Applied Basic Research Foundation (2026A1515060001, 2024A1515012003 and 2023A1515110718). Min Yang was supported by the GuangDong Basic and Applied Basic Research Foundation (2025B1515020032) and the Shenzhen Science and Technology Innovation Program (KQTD20190929172835662). We are grateful to the Shenzhen Infrastructure for Synthetic Biology for providing instrument support and technical assistance.

