## Supplementary Information for "Decoding Prokaryotic Whole Genomes with a Product-Contextualized Large Language Model"

Shiwen Ni^1^, Shuaimin Li^2^, Shijian Wang^3,4^, Wenzhi Xue^5^, Luoxi Zhang^1^, Liyang Fan^1^, Xinping Bi^2^, Yitai Li^2^, Chengguang Gan^6^, Jiarui Jin^3^, Yuan Lu^3^, Ahmadreza Argha^7^, Hamid Alinejad-Rokny^7^, Tong Si^5^, Min Yang^1,2,5^
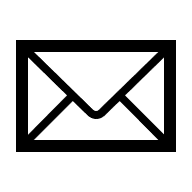
, Teng Wang^5^
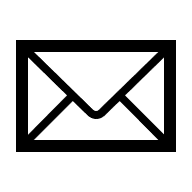

^1^Artificial Intelligence Research Institute, Shenzhen University of Advanced Technology, Shenzhen, China

^2^Shenzhen Key Laboratory for High Performance Data Mining, Shenzhen Institutes of Advanced Technology, Chinese Academy of Sciences, Shenzhen, China

^3^Xiaohongshu Inc., Shanghai, China

^4^School of Computer Science and Engineering, Southeast University, Nanjing, China

^5^State Key Laboratory of Quantitative Synthetic Biology, Shenzhen Institute of Synthetic Biology, Shenzhen Institutes of Advanced Technology, Chinese Academy of Sciences, Shenzhen, China

^6^Graduate School of Environment and Information Sciences, Yokohama National University, Yokohama, Japan

^7^School of Biomedical Engineering, UNSW Sydney, Sydney, Australia

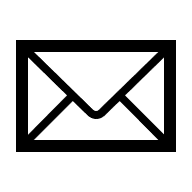

**Supplementary Figures**

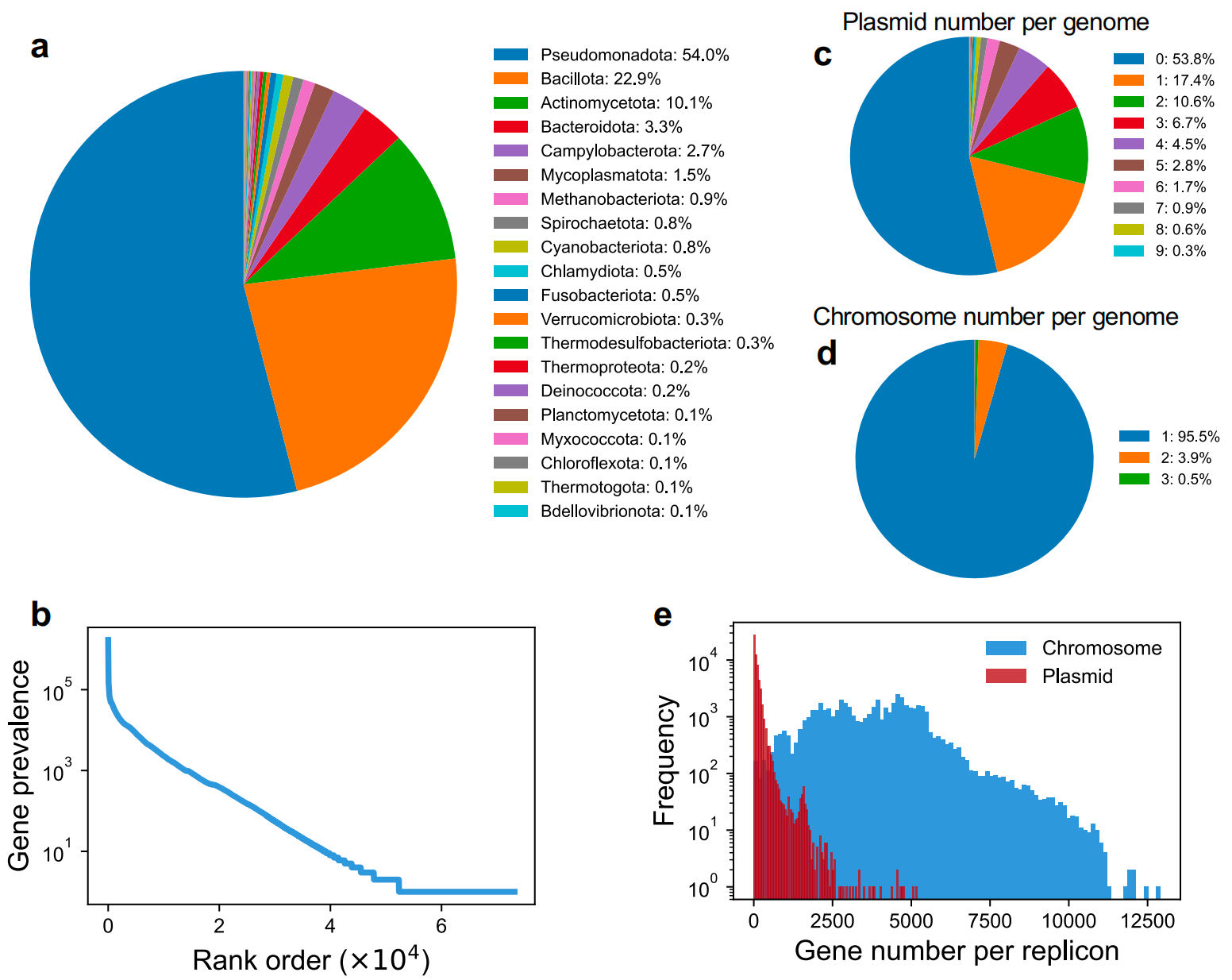

**Supplementary Figure S1| Statistical analysis of prokaryotic gene product corpus derived from NCBI RefSeq annotated genomes.** **(a)** Taxonomic distribution of the retrieved prokaryotic genomes. *Pseudomonadota* constituted the predominant phylum, accounting for the majority of sequenced genomes. **(b)** Frequency-rank distribution of gene products in the corpus follows a power-law relationship. **(c)** Distribution of plasmid number per genome. Approximately 50% of genomes harbor at least one plasmid. **(d)** Distribution of chromosome number per genome. The vast majority of genomes contained a single chromosome. **(e)** Distributions of gene number per replicon. Chromosomes and plasmids are represented by different colors.

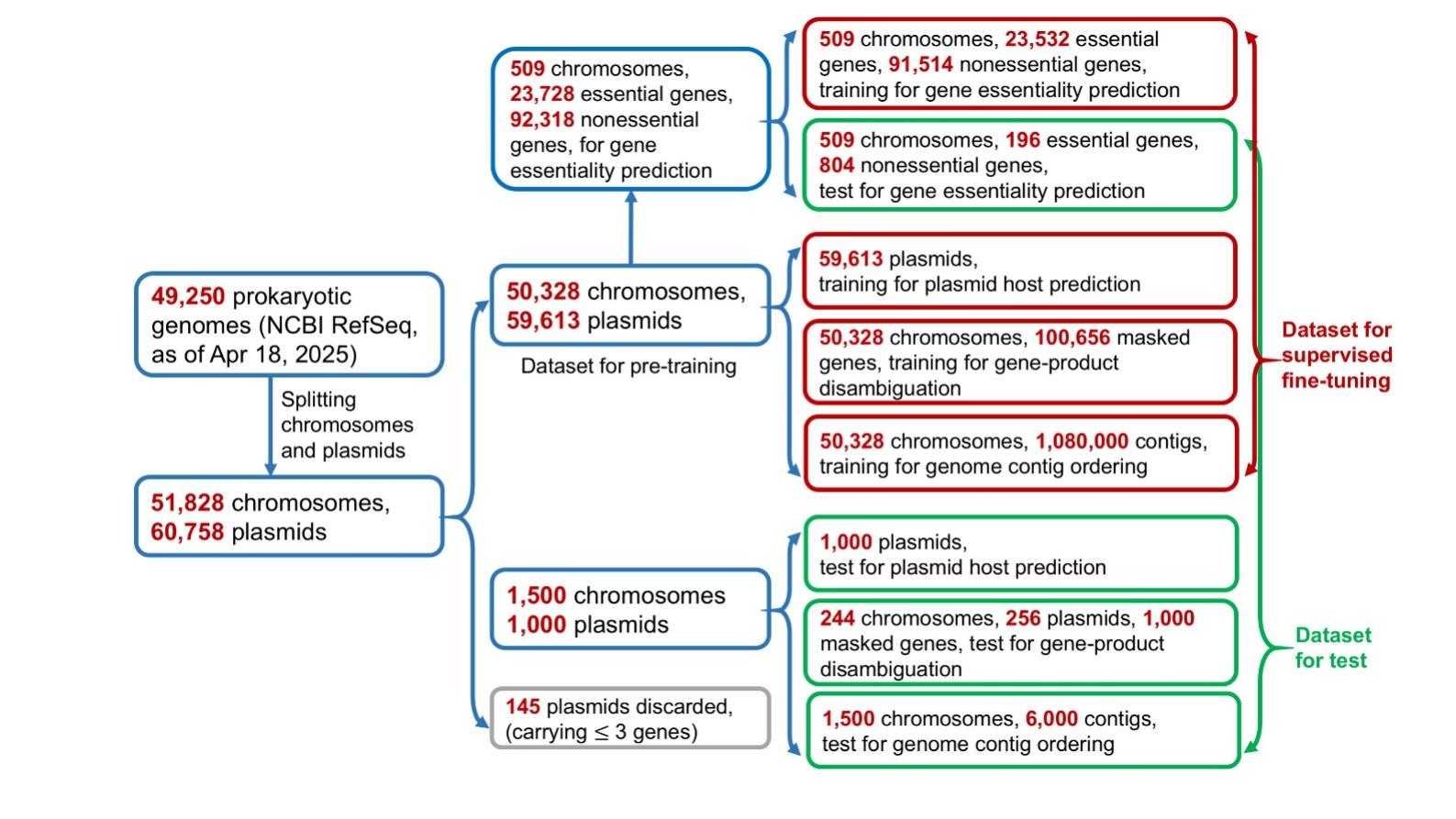

**Supplementary Figure S2| Utilization of genomic data for continuous pretraining and fine-tuning.** We compiled a dataset comprising 49,250 complete prokaryotic genomes from the NCBI RefSeq database. For genomes containing multiple replicons, we partitioned them into distinct "paragraphs", yielding 51,828 chromosomal and 60,758 plasmid segments. Of these, 50,328 chromosomes and 59,613 plasmids were utilized for continuous pre-training. These plasmids and their corresponding host information were used to fine-tune GenSyntax for plasmid host identification. Additionally, 100,656 masked genes randomly selected from the 50,328 chromosomes were used to fine-tune GenSyntax for gene-product disambiguation. We further fragmented the 50,328 complete chromosomes into 1,080,000 contigs and randomized their order to fine-tune GenSyntax for genome contig ordering. For gene essentiality prediction, essential genes from the Database of Essential Genes were incorporated into fine-tuning process. The remaining 1,500 chromosomes and 1,000 plasmids, which were not included in pre-training and fine-tuning, served as the test dataset for plasmid host prediction, gene-product disambiguation and genome contig ordering.

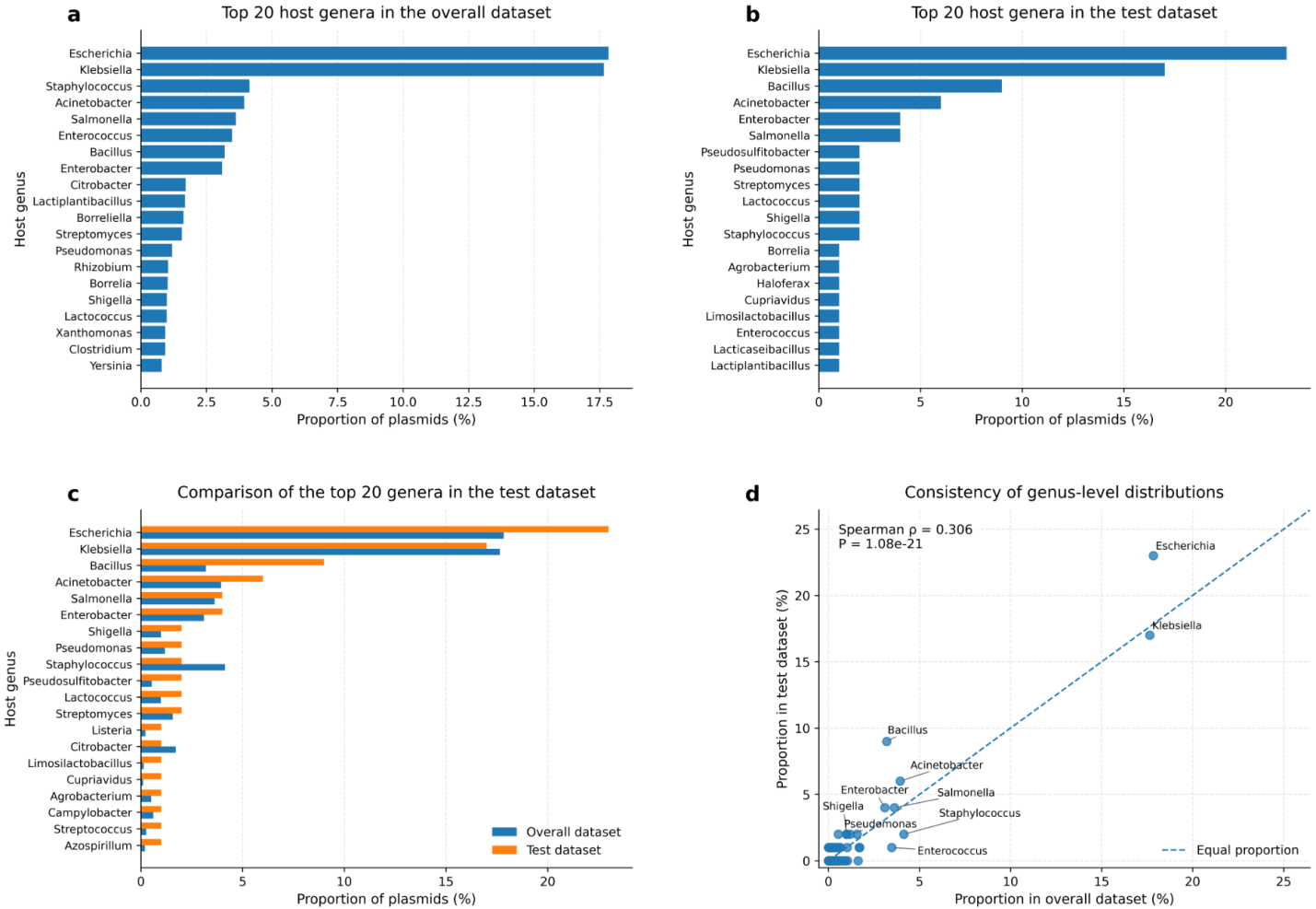

**Supplementary Figure S3| Taxonomic composition of the plasmid host prediction datasets. (a)** Distribution of the 20 most abundant host genera in the overall plasmid host prediction dataset. (**b)** Distribution of the 20 most abundant host genera in the test dataset. **(c)** Comparison of the relative proportions of the 20 most abundant genera between the overall and test datasets. (**d)** Consistency of genus-level taxonomic distributions between the overall and test datasets. Each point represents one host genus, and the dashed diagonal indicates identical proportions in the two datasets. Spearman’s rank correlation was used to quantify distributional consistency.

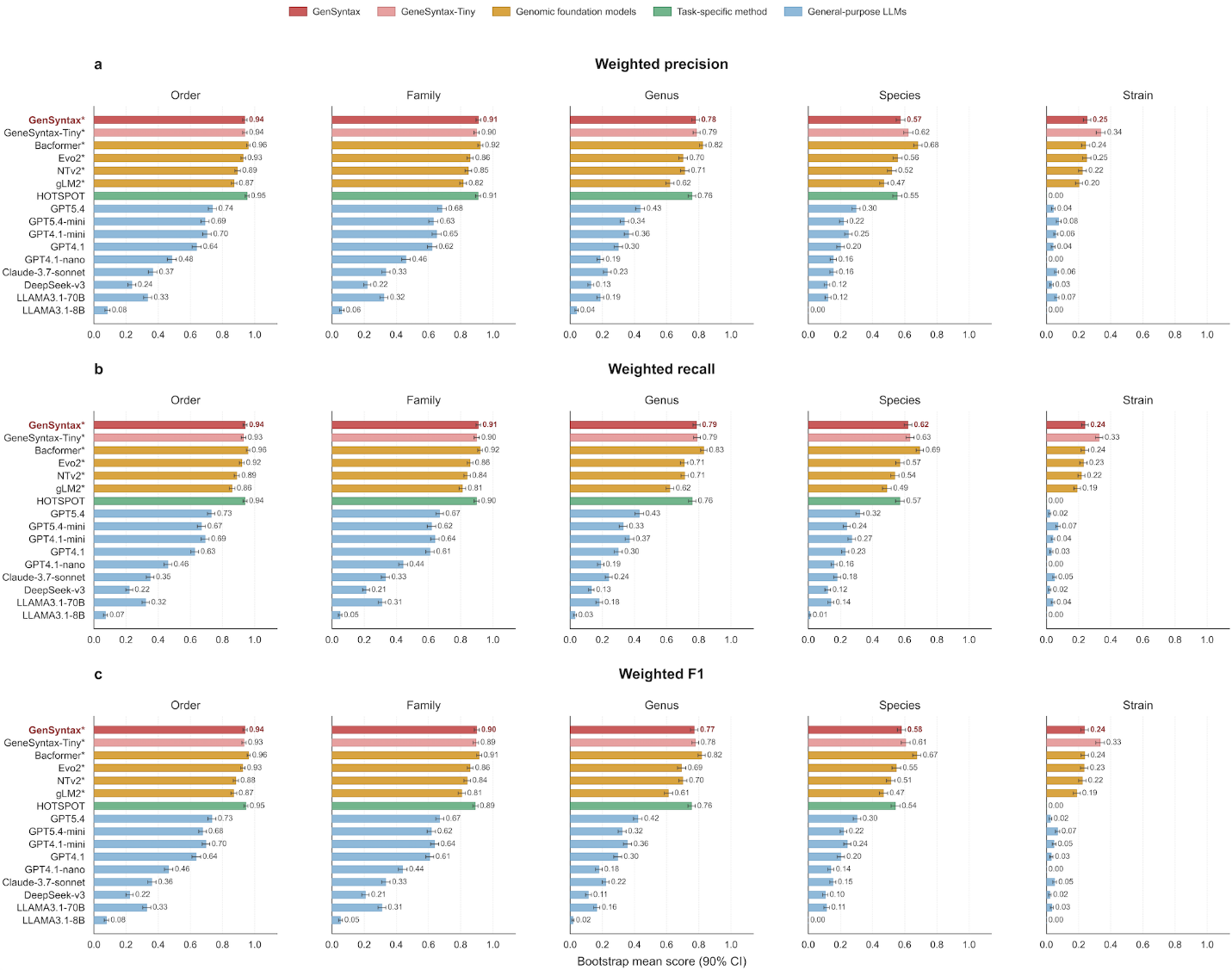

**Supplementary Figure S4 | The weighted precision, recall and F1 scores of plasmid host prediction at the order, family, genus, species and strain levels.** Performance metrics were estimated by bootstrap resampling, with points representing mean scores and error bars representing 90% confidence intervals from 100 bootstrap replicates of the test set. Comparisons included general-purpose large language models, representative genome foundation models, GenSyntax-Tiny and HOTSPOT, a domain-specific host prediction method.

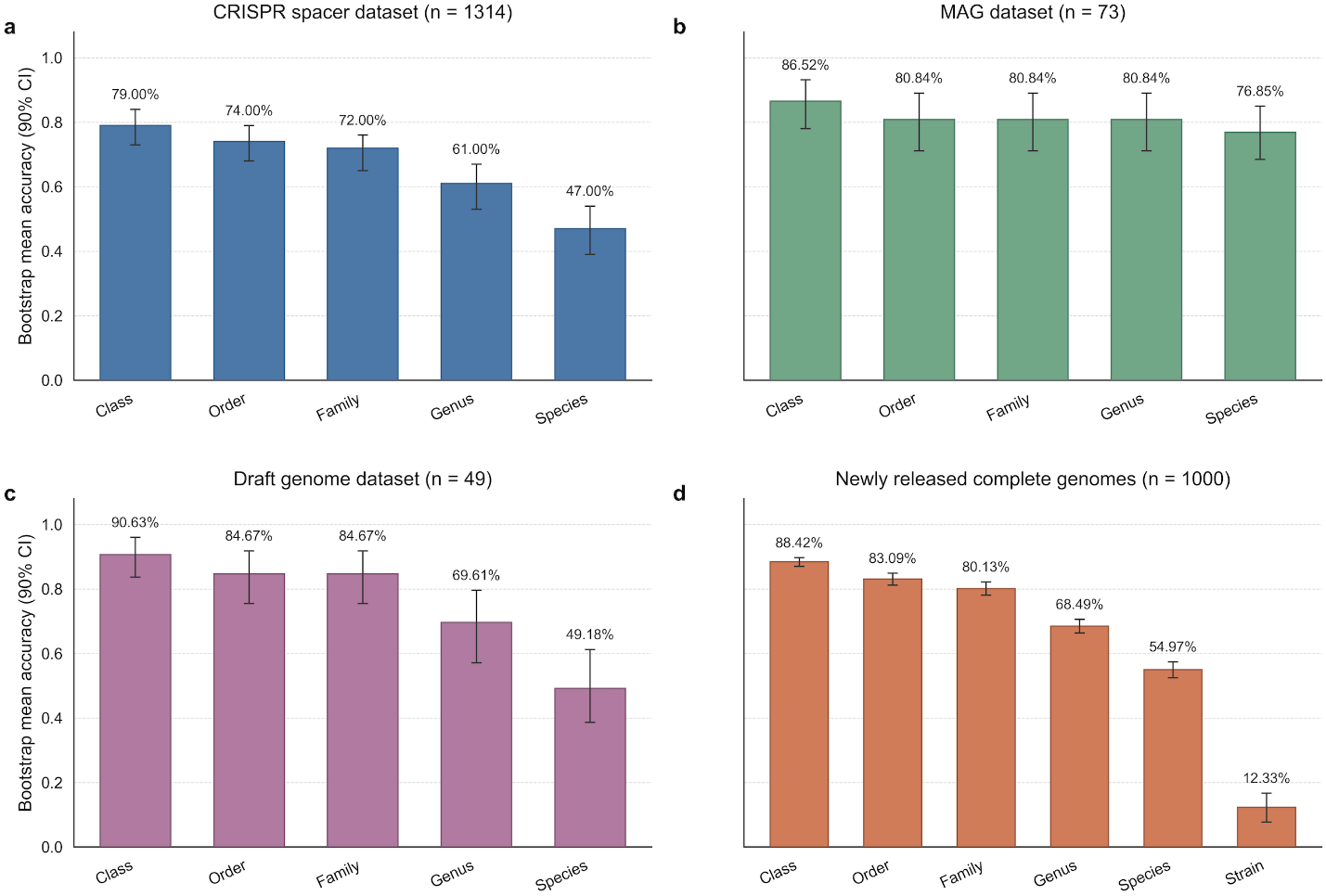

**Supplementary Figure S5| Plasmid host prediction performance evaluated on four independent external datasets.** GenSyntax was evaluated on four external validation datasets representing distinct genomic contexts. Error bars represent the mean performance and 90% confidence interval calculated from bootstrap resampling of the test set. **(a)** Complete plasmids recovered from human gut metagenomes in IMG/PR, with host assignments enabled by CRISPR spacer matches at the species level and plasmid genes independently annotated using Bakta. **(b)** Plasmids identified from metagenome-assembled genomes (MAGs). **(c)** Plasmids recovered from draft genome assemblies. (d) Plasmids from newly released complete genome assemblies. The datasets in panels (b–d) were collected from genome records released after the GenSyntax pre-training cutoff and were not used during model pre-training and task-specific fine-tuning, enabling assessment of temporal and cross-dataset generalization.
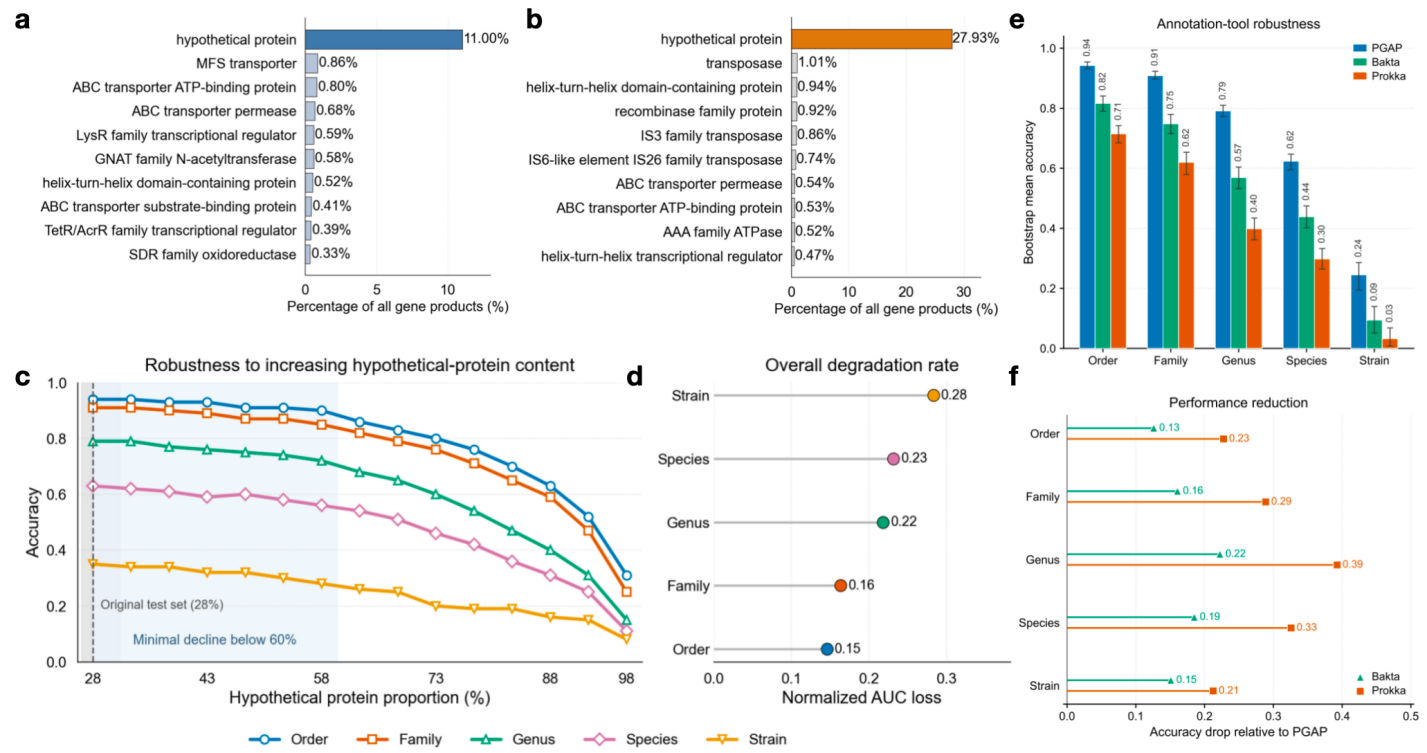

**Supplementary Figure S6| Evaluation of GenSyntax’s robustness to annotation quality.** **(a)** Top 10 most frequent gene products in the full pretraining corpus. **(b)** Top 10 most frequent gene products in the test set for plasmid host prediction. **(c)** Accuracy of GenSyntax across taxonomic ranks as the proportion of hypothetical proteins in the test set is progressively increased. The dashed line marks the original test-set proportion (28%). **(d)** Overall performance degradation across taxonomic ranks, quantified as normalized AUC loss over the full range of hypothetical-protein perturbation. **(e)** Accuracy of GenSyntax under three annotation pipelines (PGAP, Prokka and Bakta) in plasmid host prediction. **(f)** Accuracy reduction under Prokka and Bakta relative to PGAP across taxonomic ranks.

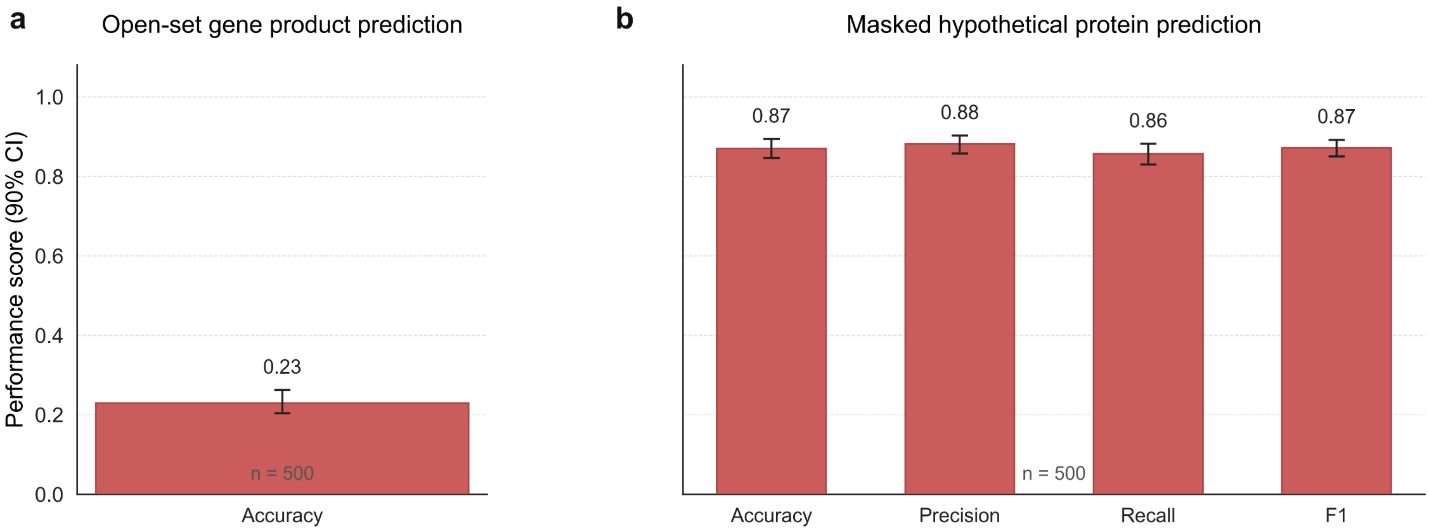

**Supplementary Figure S7| Evaluation of GenSyntax for open-domain gene function prediction and recognition of hypothetical proteins. (a)** Open-set gene function prediction, in which GenSyntax directly generates gene product annotations without being restricted to a predefined set of candidate options. Performance was evaluated by comparing the generated annotations with the ground-truth product descriptions. **(b)** Hypothetical protein recognition, in which gene product descriptors from both functionally annotated genes and genes annotated as “hypothetical protein” were randomly masked. GenSyntax was then evaluated on its ability to predict whether the masked product belonged to a hypothetical protein or a functionally annotated gene based on genomic context. Performance was assessed by comparing predicted annotation categories with the original product annotations. Error bars represent the mean performance and 90% confidence interval calculated from bootstrap resampling of the test set.

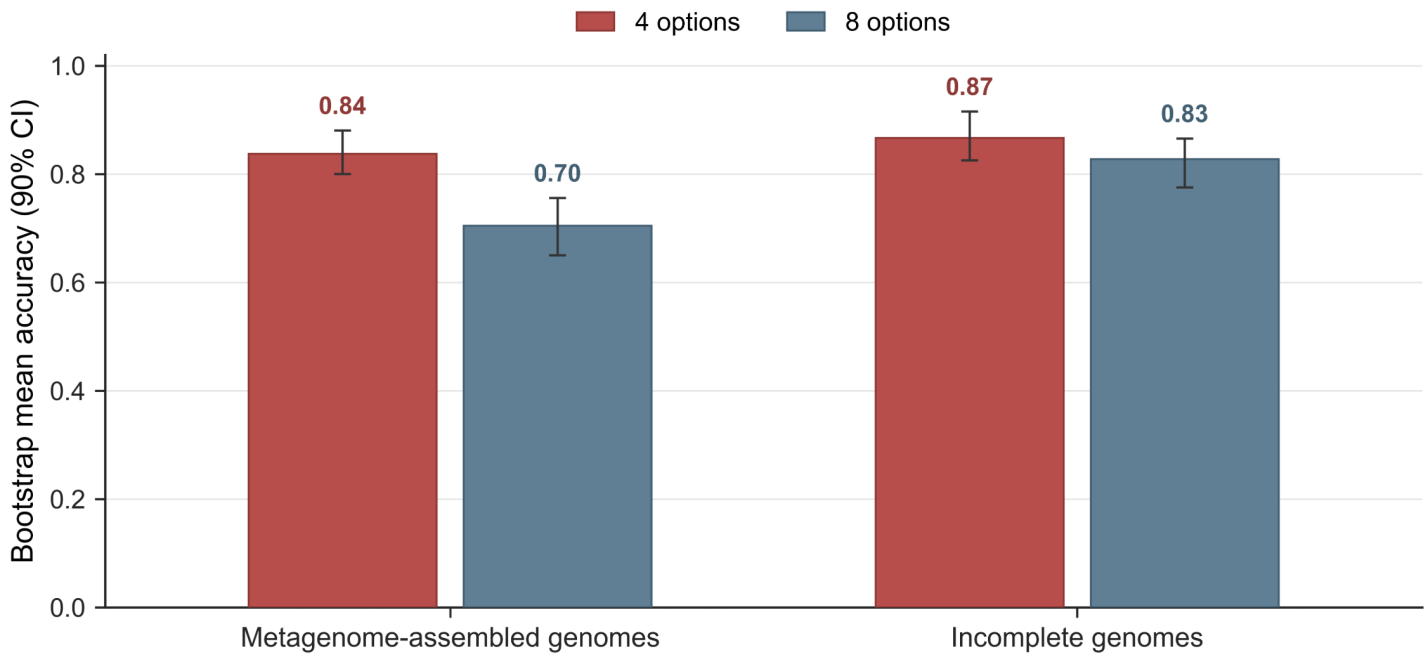

**Supplementary Figure S8| Evaluation of GenSyntax for gene function prediction in post-training draft genomes and metagenome-assembled genomes.** GenSyntax was evaluated on draft genomes and metagenome-assembled genomes (MAGs) released after the model training period, providing an independent assessment of its generalizability beyond the reference genomes used during model development. Gene function prediction performance was evaluated using the same task formulation and evaluation criteria as the original benchmark. Error bars represent the mean performance and 90% confidence interval calculated from bootstrap resampling of the test set.

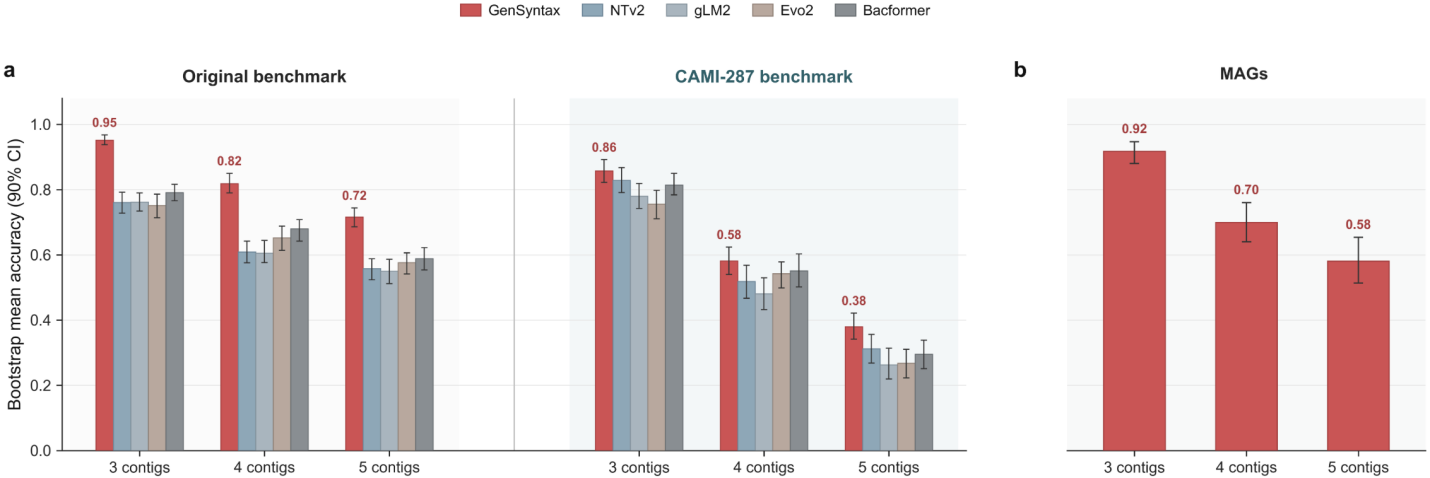

**Supplementary Figure S9 | Evaluation of GenSyntax generalization for genome contig ordering across independent genomic datasets. (a)** Performance comparison of GenSyntax and four representative genomic language models, including NTv2, gLM2, Evo2, and Bacformer, on the original contig-ordering benchmark and an independent CAMI-derived dataset. The CAMI-287 benchmark was constructed from genomes excluded from the GenSyntax training corpus to provide an external evaluation of model generalization. **(b)** Evaluation of GenSyntax on metagenome-assembled genomes (MAGs) released to RefSeq after the GenSyntax training period. These MAGs were not available during model development and were used to assess the ability of GenSyntax to recover genome organization in externally generated, non-reference genomic datasets. Error bars represent the mean performance and 90% confidence interval calculated from bootstrap resampling of the test set.

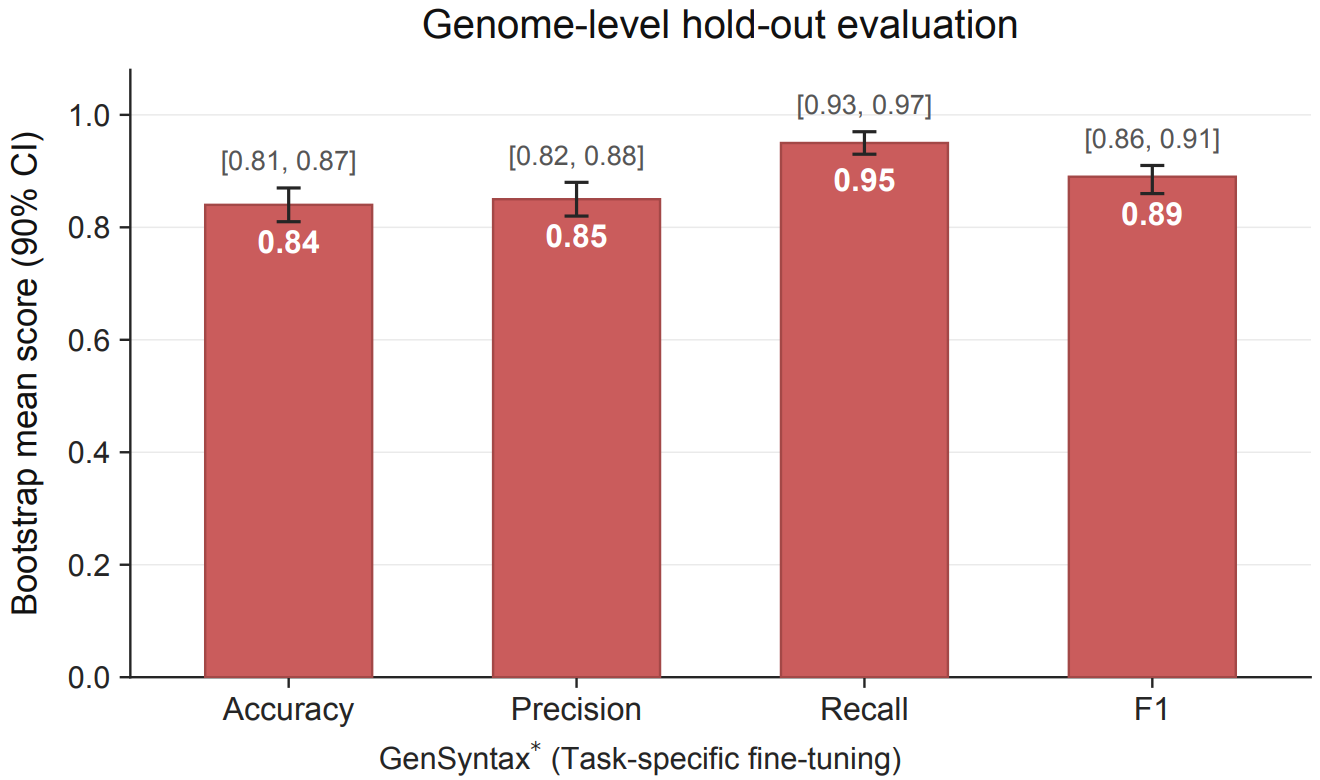

**Supplementary Figure S10 | Evaluation of GenSyntax gene essentiality prediction under a genome-level hold-out strategy.** GenSyntax performance was reassessed using a stricter genome-level data partitioning scheme, in which all genes from the same genome were assigned exclusively to either the training or test set. The model maintained robust gene essentiality prediction performance under this more stringent evaluation setting, demonstrating that its predictions generalize beyond genomic contexts encountered during model training.

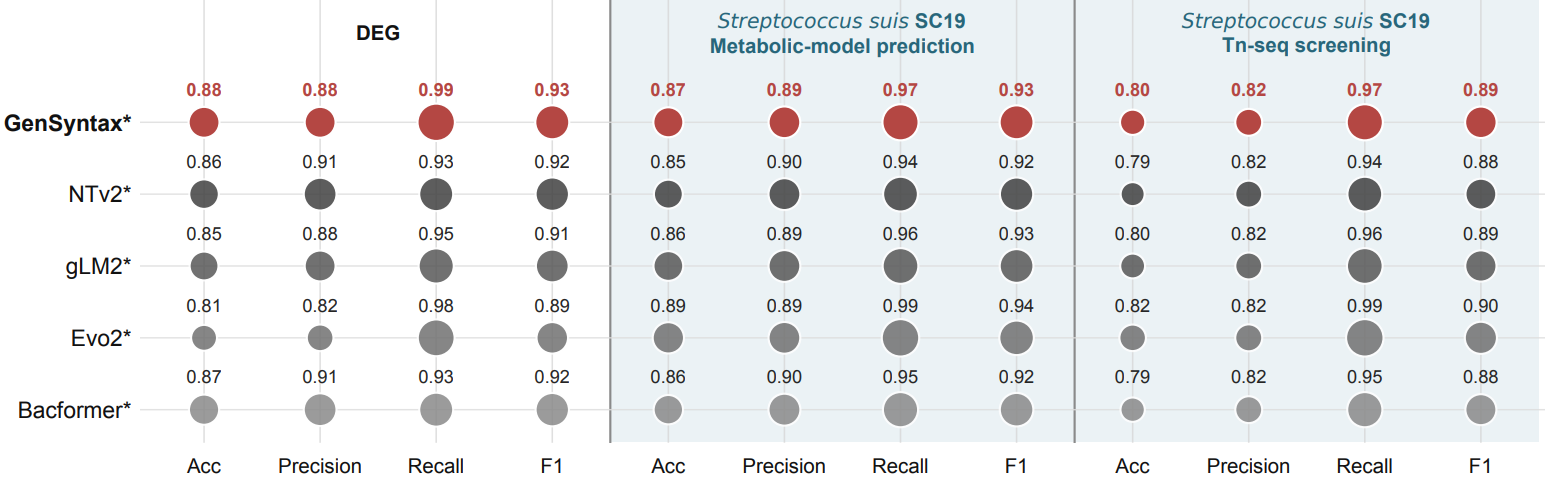

**Supplementary Figure S11| Generalization performance of GenSyntax for gene essentiality prediction in an independent benchmark.** GenSyntax was evaluated on the original Database of Essential Genes (DEG)-derived benchmark and two independent external essentiality datasets based on complementary computational and experimental approaches: genome-scale metabolic model predictions and transposon insertion sequencing (Tn-seq) screening. These external benchmarks were not used during model training and provide an assessment of whether GenSyntax captures transferable signals associated with gene essentiality beyond the original DEG annotations. The results demonstrate robust performance across datasets generated from distinct essentiality inference strategies.

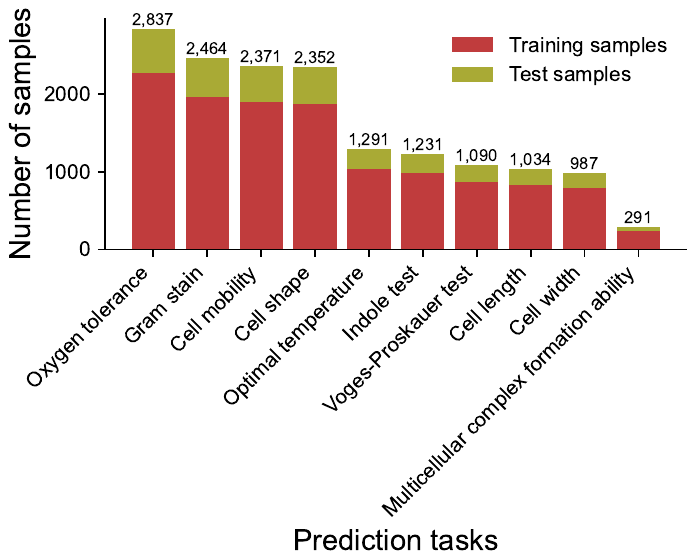

**Supplementary Figure S12| The sample sizes for training and test sets for each phenotype prediction.** Ten microbial phenotype datasets were retrieved from the BacDive database through its Advanced Search interface. Species names were mapped to genome accessions in RefSeq to establish integrated genotype-phenotype datasets for all ten phenotypes. Each dataset was randomly partitioned into training and test sets at a 4:1 ratio.

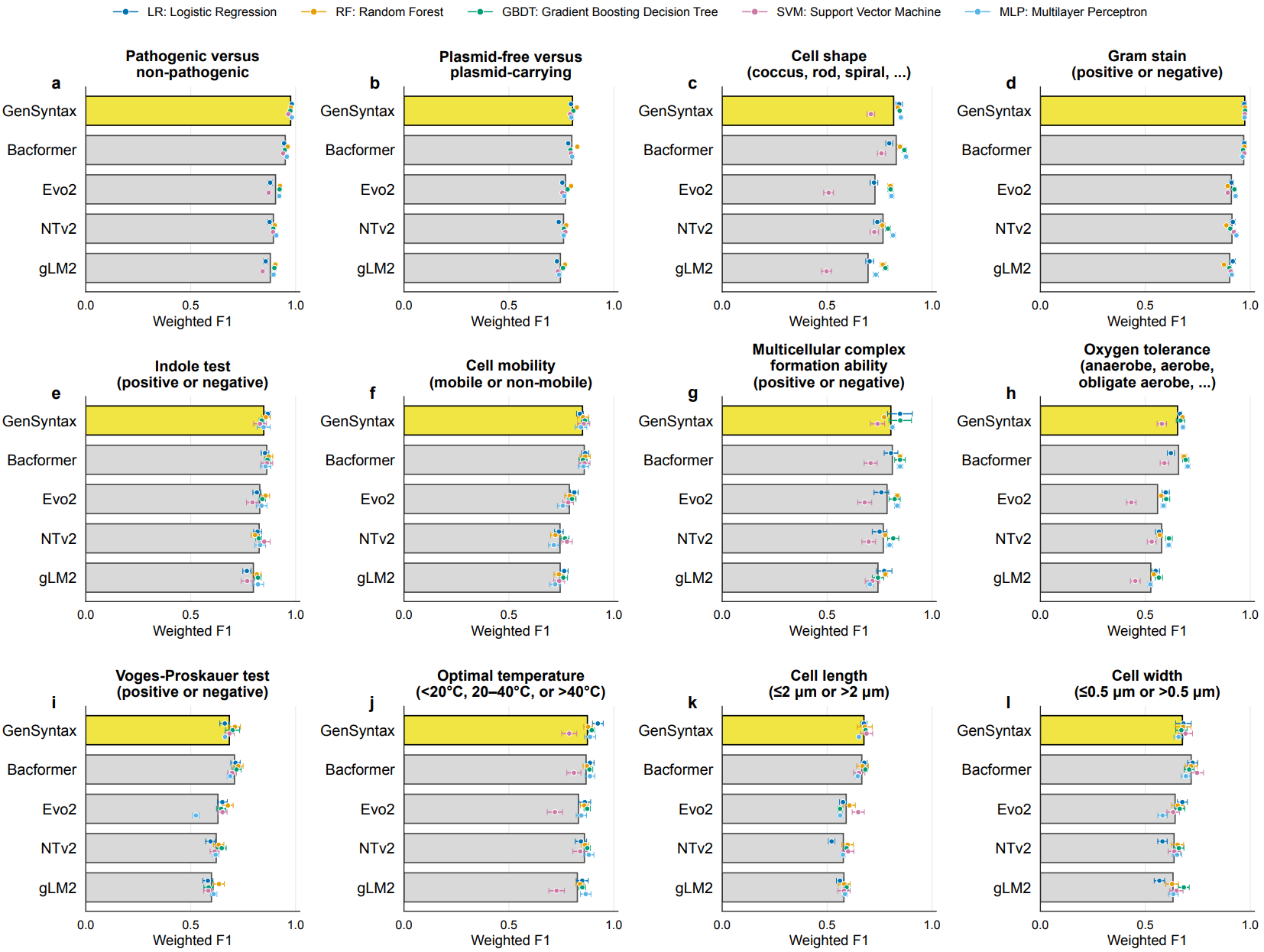

**Supplementary Figure S13| Benchmarking genome language model embeddings for microbial phenotype prediction.** Chromosome-level representations generated by GenSyntax and four baseline genome language models, including Bacformer, NT-v2, Evo2, and gLM2, were used to predict a broad spectrum of microbial traits compiled from the BacDive database and other curated phenotype labels. Five machine learning classifiers were used for prediction: Logistic Regression (LR), Random Forest (RF), Support Vector Machine (SVM), Gradient Boosting Decision Tree (GBDT), and Multilayer Perceptron (MLP). Each panel shows prediction accuracies for a different phenotype or genomic trait: **(a)** distinguishing pathogenic from non-pathogenic strains, **(b)** identifying whether a genome carries plasmids, **(c)** classifying cell shape, such as coccus, rod, and spiral, **(d)** Gram staining reaction, positive versus negative, **(e)** indole test outcome, positive versus negative, **(f)** cell motility, mobile versus non-mobile, **(g)** ability to form multicellular complexes, positive versus negative, **(h)** oxygen tolerance, such as anaerobe, aerobe, and facultative anaerobe, **(i)** Voges–Proskauer test outcome, positive versus negative, **(j)** preferred growth temperature, <20 °C, 20–40 °C, or >40 °C, **(k)** average cell length, ≤2 µm versus >2 µm, and **(l)** average cell width, ≤0.5 µm versus >0.5 µm. Bar lengths represent the mean F1 across classifiers for each genome language model, and colored dots indicate the accuracies obtained by individual classifiers. Each dot represents the mean ± standard deviation of three replicates.

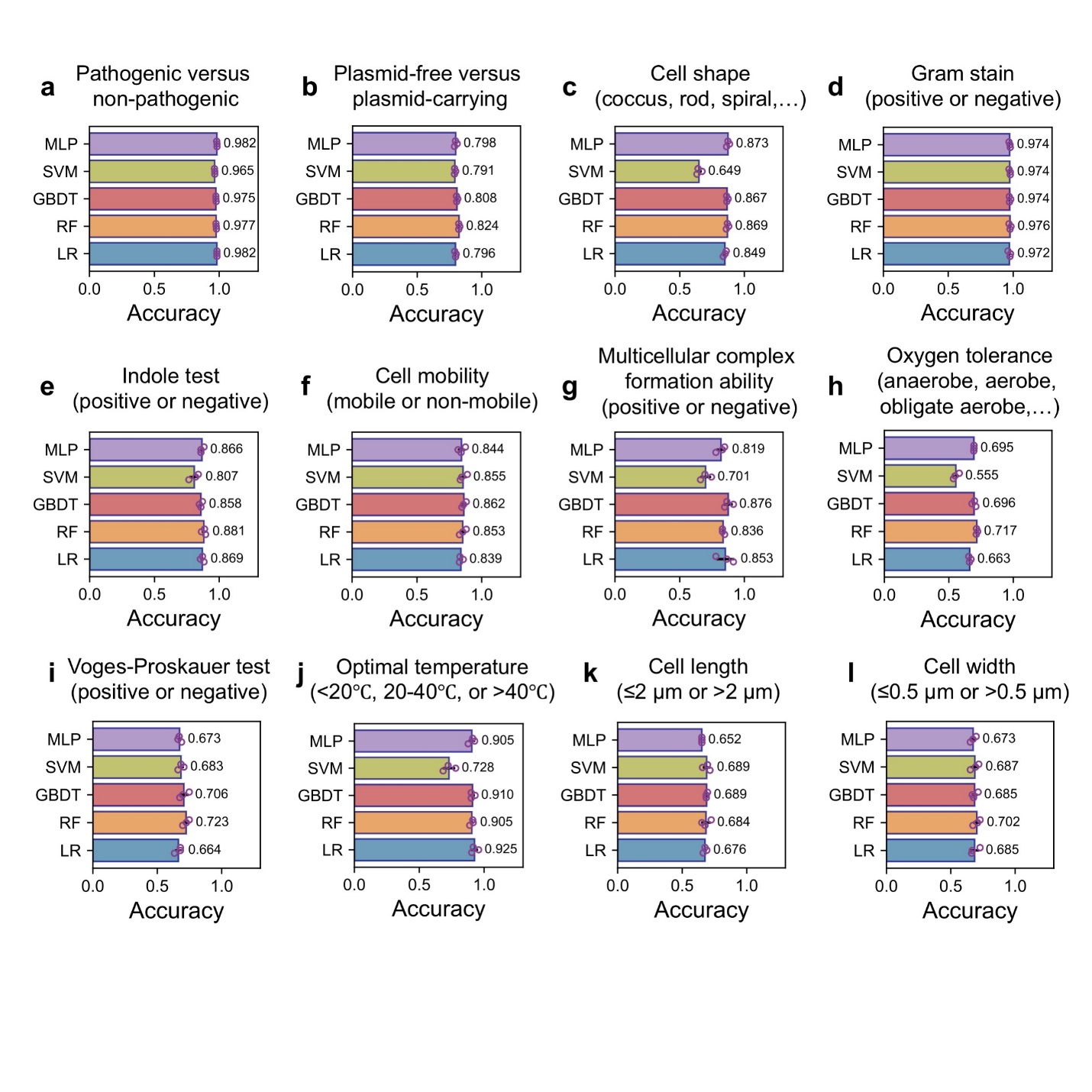

**Supplementary Figure 14| The accuracies of GenSyntax prediction of diverse microbial phenotypes.** We used chromosome representations generated by GenSyntax to predict a broad spectrum of microbial traits compiled from the BacDive database. Five machine learning classifiers were used for predicitons: Logistic Regression (LR), Random Forest (RF), Support Vector Machine (SVM), Gradient Boosting Decision Tree (GBDT), and Multilayer Perceptron (MLP). Each panel illustrates prediction accuracies for a different phenotype: **(a)** distinguishing pathogenic from non-pathogenic strains, **(b)** identifying whether a genome carries plasmids, **(c)** classifying cell shape (e.g., coccus, rod, spiral), **(d)** Gram staining reaction (positive vs. negative), **(e)** indole test outcome (positive vs. negative), **(f)** cell motility (mobile vs. nonmobile), **(g)** ability to form multicellular complexes (positive vs. negative), **(h)** oxygen tolerance (e.g., anaerobe, aerobe, facultative anaerobe), **(i)** Voges–Proskauer test outcome (positive vs. negative), **(j)** preferred growth temperature (<20 °C, 20–40 °C, or >40 °C), **(k)** average cell length (≤2 µm vs. >2 µm), and **(l)** average cell width (≤0.5 µm vs. >0.5 µm). Data were presented as mean ± standard deviation of three replicates.

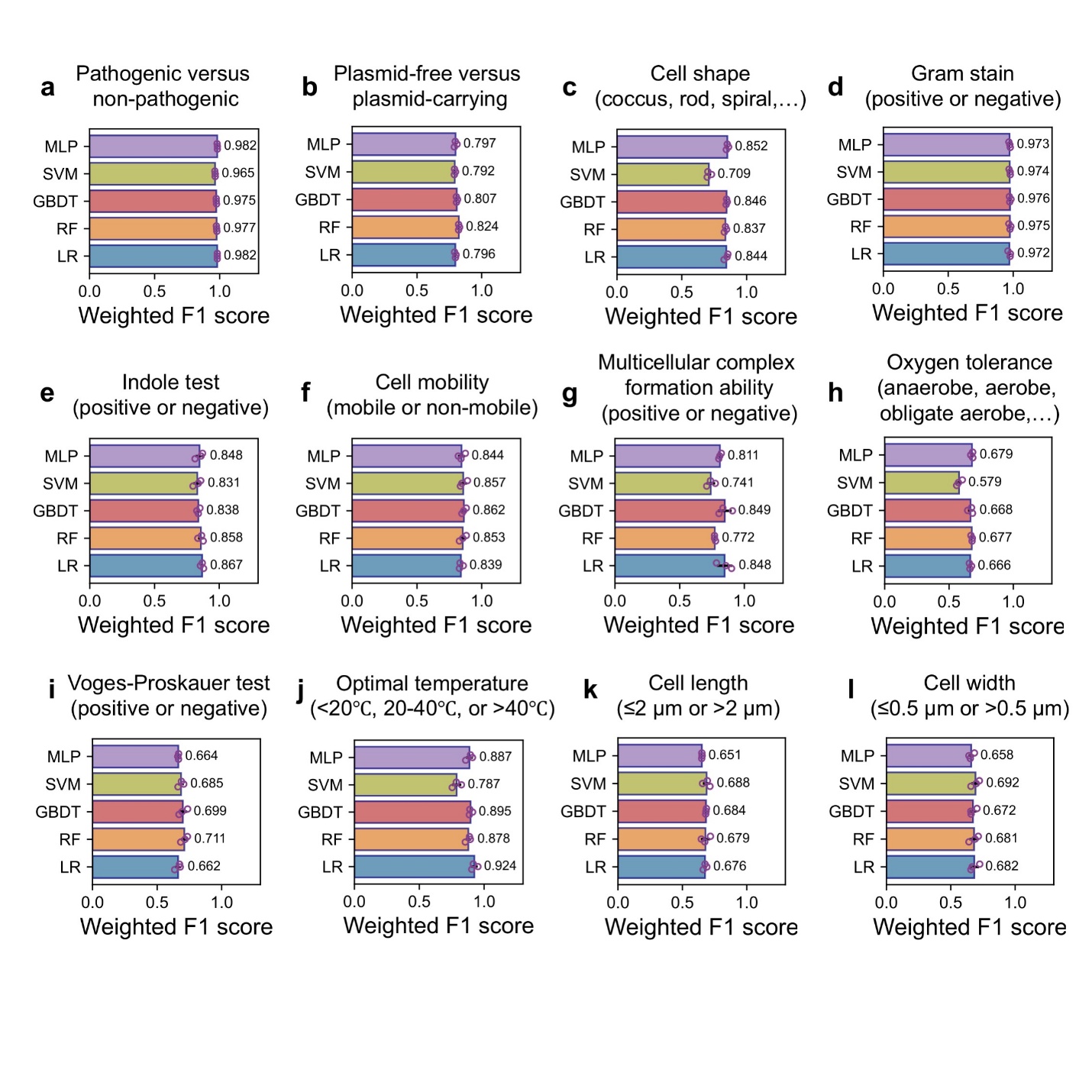

**Supplementary Figure S15| The weighted F1 scores of GenSyntax prediction of diverse microbial phenotypes.** We used five machine learning classifiers for predicitons: Logistic Regression (LR), Random Forest (RF), Support Vector Machine (SVM), Gradient Boosting Decision Tree (GBDT), and Multilayer Perceptron (MLP). Each panel illustrates the F1 scores for a different phenotype: **(a)** distinguishing pathogenic from non-pathogenic strains, **(b)** identifying whether a genome carries plasmids, **(c)** classifying cell shape (e.g., coccus, rod, spiral), **(d)** Gram stain reaction (positive vs. negative), **(e)** indole test outcome (positive vs. negative), **(f)** cell motility (mobile vs. nonmobile), **(g)** ability to form multicellular complexes (positive vs. negative), **(h)** oxygen tolerance (eg., anaerobe, aerobe, facultative anaerobe), **(i)** Voges–Proskauer test outcome (positive vs. negative), **(j)** preferred growth temperature (<20 °C, 20–40 °C, or >40 °C), **(k)** average cell length (≤2 µm vs. >2 µm), and **(l)** average cell width (≤0.5 µm vs. >0.5 µm). Data were presented as mean ± standard deviation of three replicates.

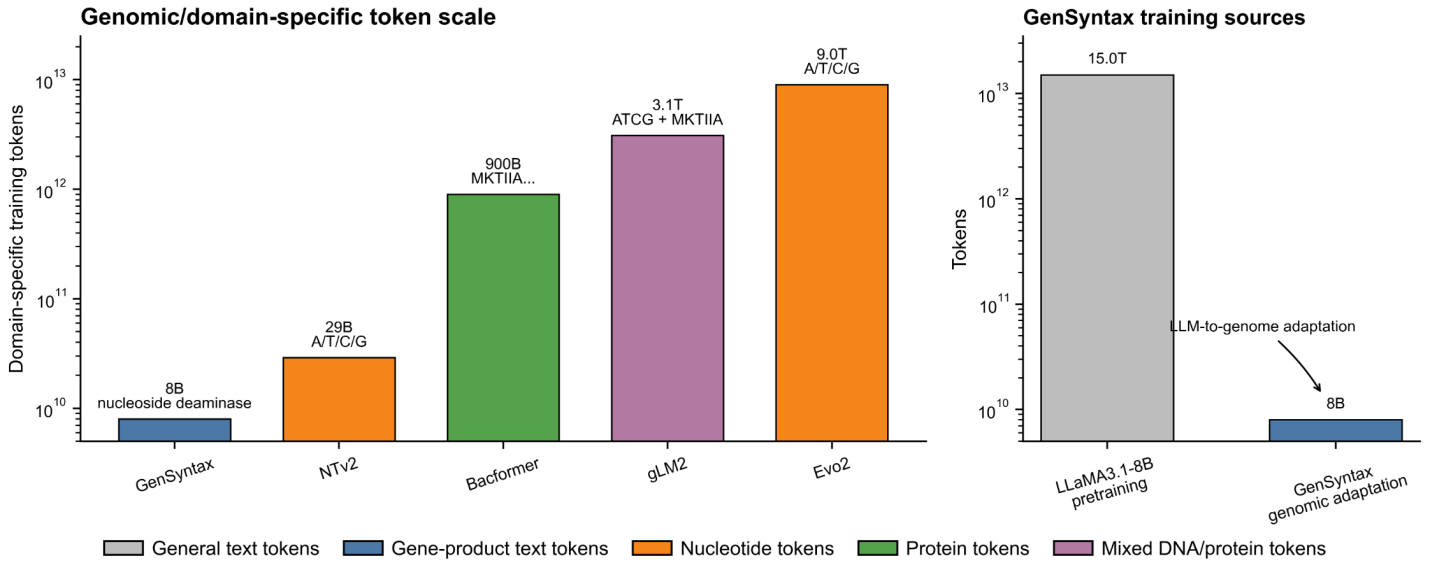

**Supplementary Figure S16| Comparison of genomic or domain-specific token scales across GenSyntax and representative genomic foundation models, together with the contribution of LLaMA3.1-8B pretraining and GenSyntax genomic adaptation.** Token counts are shown on a log scale and colored by input modality.

| Functional_category | Syn3A | | Syn3B | | GenSyntax_0.5 | | GenSyntax_0.4 | | GenSyntax_0.3 | | GenSyntax_0.2 | |
| --- | --- | --- | --- | --- | --- | --- | --- | --- | --- | --- | --- | --- |
|  | kept | deleted | kept | deleted | kept | deleted | kept | deleted | kept | deleted | kept | deleted |
| Glucose transport and glycolysis | 10 | 2 | 10 | 2 | 7 | 5 | 7 | 5 | 7 | 5 | 11 | 1 |
| Ribosome biogenesis | 6 | 1 | 6 | 1 | 5 | 2 | 7 | 0 | 6 | 1 | 7 | 0 |
| Protein export | 9 | 0 | 9 | 0 | 8 | 1 | 9 | 0 | 9 | 0 | 9 | 0 |
| Transcription | 10 | 6 | 10 | 6 | 5 | 11 | 7 | 9 | 7 | 9 | 9 | 7 |
| RNA metabolism | 13 | 1 | 13 | 1 | 1 | 13 | 4 | 10 | 5 | 9 | 9 | 5 |
| DNA topology | 5 | 0 | 5 | 0 | 4 | 1 | 4 | 1 | 5 | 0 | 5 | 0 |
| Chromosome segregation | 2 | 0 | 2 | 0 | 0 | 2 | 1 | 1 | 2 | 0 | 2 | 0 |
| DNA metabolism | 5 | 2 | 5 | 2 | 1 | 6 | 4 | 3 | 2 | 5 | 7 | 0 |
| Protein folding | 3 | 1 | 3 | 1 | 2 | 2 | 3 | 1 | 3 | 1 | 4 | 0 |
| Translation | 81 | 1 | 80 | 2 | 73 | 9 | 78 | 4 | 79 | 3 | 82 | 0 |
| RNA (rRNAs, tRNAs, small RNAs) | 37 | 2 | 38 | 1 | 8 | 31 | 22 | 17 | 35 | 4 | 38 | 1 |
| DNA replication | 11 | 3 | 11 | 3 | 11 | 3 | 13 | 1 | 12 | 2 | 13 | 1 |
| Lipid salvage and biogenesis | 7 | 0 | 7 | 0 | 5 | 2 | 7 | 0 | 5 | 2 | 7 | 0 |
| Cofactor transport and salvage | 4 | 4 | 5 | 3 | 2 | 6 | 3 | 5 | 3 | 5 | 7 | 1 |
| rRNA modification | 7 | 1 | 7 | 1 | 0 | 8 | 1 | 7 | 0 | 8 | 3 | 5 |
| tRNA modification | 9 | 2 | 9 | 2 | 4 | 7 | 5 | 6 | 7 | 4 | 10 | 1 |
| Efflux | 1 | 2 | 1 | 2 | 0 | 3 | 0 | 3 | 0 | 3 | 0 | 3 |
| Nucleotide salvage | 16 | 2 | 16 | 2 | 5 | 13 | 10 | 8 | 9 | 9 | 15 | 3 |
| DNA repair | 4 | 2 | 4 | 2 | 0 | 6 | 0 | 6 | 1 | 5 | 5 | 1 |
| Metabolic processes | 65 | 86 | 65 | 86 | 25 | 126 | 36 | 115 | 45 | 106 | 88 | 63 |
| Membrane transport | 35 | 51 | 36 | 50 | 4 | 82 | 4 | 82 | 10 | 76 | 59 | 27 |
| Redox homeostasis | 5 | 0 | 5 | 0 | 0 | 5 | 3 | 2 | 2 | 3 | 5 | 0 |
| Proteolysis | 11 | 18 | 11 | 18 | 2 | 27 | 3 | 26 | 5 | 24 | 13 | 16 |
| Regulation | 0 | 1 | 0 | 1 | 0 | 1 | 0 | 1 | 0 | 1 | 0 | 1 |
| Unassigned | 113 | 156 | 113 | 156 | 17 | 252 | 23 | 246 | 28 | 241 | 94 | 175 |
| Cell division | 4 | 0 | 4 | 0 | 1 | 3 | 2 | 2 | 3 | 1 | 3 | 1 |
| Lipoprotein | 11 | 48 | 11 | 48 | 1 | 58 | 1 | 58 | 3 | 56 | 33 | 26 |
| Transport and catabolism of nonglucose carbon sources | 4 | 2 | 4 | 2 | 1 | 5 | 3 | 3 | 2 | 4 | 4 | 2 |
| Acylglycerol breakdown | 1 | 1 | 1 | 1 | 0 | 2 | 0 | 2 | 0 | 2 | 0 | 2 |
| Mobile elements and DNA restriction | 0 | 17 | 2 | 15 | 0 | 17 | 0 | 17 | 1 | 16 | 3 | 14 |
| Total | 489 | 412 | 493 | 408 | 192 | 709 | 260 | 641 | 296 | 605 | 545 | 356 |

**Supplementary Table S1. Functional composition of the minimal genomes Syn3A and Syn3B and minimal genomes predicted by GenSyntax at different essentiality thresholds.** Genes were grouped by functional category and classified as retained (kept) or removed (deleted). GenSyntax results are reported at essentiality probability thresholds of 0.5, 0.4, 0.3 and 0.2.
